# The myeloid cell-driven transdifferentiation of endothelial cells into pericytes promotes the restoration of BBB function and brain self-repair after stroke

**DOI:** 10.1101/2023.12.07.570712

**Authors:** Tingbo Li, Ling Yang, Jiaqi Tu, Yufan Hao, Zhu Zhu, Yingjie Xiong, Qingzhu Gao, Lili Zhou, Guanglei Xie, Dongdong Zhang, Xuzhao Li, Yuxiao Jin, Yiyi Zhang, Bingrui Zhao, Nan Li, Xi Wang, Jie-Min Jia

## Abstract

Ischemic stroke, one of the leading causes of death in the world, is accompanied by the dysfunction of the blood-brain barrier (BBB), which aggravates neuron damage. However, the mechanisms underlying the restoration of BBB in the chronic stage after stroke remain unclear. Here, we investigated the changes in the pericyte pool and their effects on BBB function and brain self-repair after stroke. Using lineage tracing, RNA-seq and immunofluorescence staining, we detected endothelial cells (ECs) transdifferentiated into pericytes (E-pericytes) after stroke in the MCAO model. E-pericytes depletion by diphtheria toxin A (DTA) aggravated BBB leakage and exacerbated neurological deficits in the MCAO model. The myeloid cell-driven transdifferentiation of ECs into pericytes accelerated BBB restoration and brain self-repair after stroke via endothelial-mesenchymal transformation (EndoMT). Decreasing the number of E-pericytes by specific knockout of the *Tgfbr2* gene in ECs also aggravated BBB leakage and exacerbated neurological deficits. Specific-ECs overexpression of the *Tgfbr2* gene promoting E-pericytes transdifferentiation reduced BBB leakage and exerted neuroprotective effects. Our findings solve a key question about how changes in the pericyte pool after stroke affect the restoration of BBB function and brain self-repair and may offer a new approach to therapy.

## Introduction

All vertebrate creatures have evolved strategies for tissue repair after an accident or conflict and effective tissue repair is critical for survival. The liver is the only solid organ that uses regenerative mechanisms to ensure that the liver-to-bodyweight ratio is always at 100% of what is required for body homeostasis^1,2^. Vascular repair is crucial in liver regeneration because blood vessels can provide nutrients and oxygen to facilitate tissue restoration and functional recovery in the chronic stage^3^. Vascular repair is essential in skin wound healing as it promotes the formation of new blood vessels, enhances tissue regeneration, and supports the delivery of nutrients and oxygen to the wound site in larger injuries^4,5^. The liver and skin demonstrate a powerful ability for tissue self-repair, which benefits from vascular repair. As one of the most important organs, current knowledge and research on the contribution of vascular repair to brain self-repair are inadequate.

Stroke results in large-scale cell death because of ongoing ischemia and hypoxia, which remain a major cause of adult disability and death^6–8^. The definitive spontaneous repairing processes in the brain remain largely unknown after stroke and the decoding and harnessing of these processes may accelerate brain recovery which usually takes years. Until recently, such efforts mainly involved protective factors and cell transplants designed to rescue or replace a specific population of neurons, and the results have largely been disappointing in clinical outcomes^9–12^. Treatments to reduce neuronal death and limit acute damage are desirable by timely restoring blood vessel reperfusion in the cute phase^13,14^. No proven medical therapies exist for restoring cerebral blood flow (CBF) in the chronic phase. Unfortunately, the knowledge of brain-self repair by remedying vascular vessels after stroke in the chronic repair phase is still rare.

The vascular hierarchy comprises arteries, arterioles, capillaries, venules, and veins. Nutrient and gas exchange occurs primarily in the capillary beds, up to 90% of blood vessel length^15,16^. Capillaries are mainly composed of endothelial cells (ECs) and pericytes, which safeguard the functions of capillary beds^17^. ECs and pericytes respond to different reactions and fates after stroke. Pericytes rapidly exhibited apoptosis and death after stroke, nevertheless ECs tolerance ischemia and hypoxia^18^. When pericytes were lost, the functions of vascular vessels, BBB and CBF, were destroyed. BBB serves as a critical selective barrier that regulates the exchange of substances between the blood and the central nervous system (CNS), maintaining an obligatory environment for cell communications^19,20^. Effective CBF is essential for cell survival in the brain after stroke. The prevention of BBB leakage and the decrease of CBF due to pericyte loss is important for vascular and neurological functions after stroke.

The endothelium is capable of remarkable plasticity. Certain ECs in the embryo undergo hematopoietic transition, giving rise to multi-lineage hematopoietic stem and progenitor cells, which are crucial for developing the blood system^21^. ECs can undergo EndoMT, acquiring mesenchymal properties. This process is important in various physiological and pathological conditions, such as fibrosis and cancer^22^. Endocardial endothelial cells are progenitors of pericytes and vascular smooth muscle cells in the murine embryonic heart via EndoMT^23^. Protein C receptor-expressing (Procr^+^) ECs would give rise to de novo formation of ECs and pericytes in the mammary gland^24^. In situ, is there a self-repair mechanism to replenish the pericytes pool from ECs for protection functions of blood vessels after stroke in the chronic repair phase that is unknown?

In the acute and subacute phases after stroke, myeloid cells were dominant immune cells entering the brain parenchyma^25^. Most myeloid cells could not occupy the ischemic brain for a long time and faded away by releasing various factors, including pro-inflammatory, anti-inflammatory and trophic factors^26–29^. Blocking myeloid cell recruitment using anti-CCR2 antibody and *CCR2* gene knockout mice impaired long-term spontaneous behavioral recovery after stroke via reducing anti-inflammatory macrophages and angiogenesis^30–32^. TGF-β1 was predominantly co-localized with CD68^+^ activated microglia and macrophage in mice after distal middle cerebral artery occlusion (dMCAO)^33^ and TGF-β1 is important in the pathogenesis of the angiogenic response in ischemic brain tissue in patients after stroke^34^. At the same time, TGF-β is the main driver of EndoMT, which is involved in endothelial-to-pericytic transition in the murine embryonic heart and mammary gland^35^. However, it remains unclear whether myeloid cells can drive endothelial-to-pericytic transition to replenish the pericyte pool and remedy the functions of vascular vessels after stroke in the chronic phase.

The study was to verify that ECs can transdifferentiate into E-pericytes to replenish the pericyte pool in situ and assess the role of E-pericytes in the restoration of BBB function and brain self-repair. In the MCAO model, we discovered that ischemic stroke induced EC-to-pericyte transdifferentiation. Myeloid cells drive E-pericytes via the TGFb1-TGFβR2 pathway. The specific decrease in the number of E-pericytes by DTA and knockout of the *Tgfbr2* gene in ECs increased BBB leakage and impaired long-term spontaneous behavioral recovery. Increasing the number of E-pericytes via overexpression of the *Tgfbr2* gene in ECs alleviated BBB leakage and enhanced long-term spontaneous behavioral recovery. These results reveal the previously unappreciated transdifferentiation of E-pericyte to replenish the pericyte pool after, its role in BBB restoration and brain self-repair after stroke and how infiltrating myeloid cells drive E-pericyte formation. Our study uncovered a new mechanism for accelerating brain self-repair and provided a new approach to therapy after stroke.

## Results

### Changes in the pericyte pool after stroke

To study EC-to-pericyte transdifferentiation after stroke and the underlying mechanisms, we first aimed to explore changes in the pericyte pool after stroke. We used the MCAO model in mice to mimic stroke patients^26^. CD13^+^ pericytes and CD31^+^ ECs were found to be TUNEL positive on capillary in our stroke model. Moreover, only half of the pericytes survived at reperfusion 2 days (RP2D), while over 80% of ECs survived (Figure 1A). The number of CD13^+^ pericytes on capillary also decreased but then increased at RP2D and RP7D after stroke (Figure 1B-C). Flow cytometry also revealed that the number of pericytes significantly decreased, from 100% on the contralateral side to approximately 50% on the ipsilateral side at RP2D, but gradually increased following reperfusion after stroke (Figure 1D-E). Overall, there was a 68.1% increase in the pericyte pool from RP2D to RP34D and a 40.5% increase in total pericytes at RP34D (Figure 1E).

**Figure 1.**
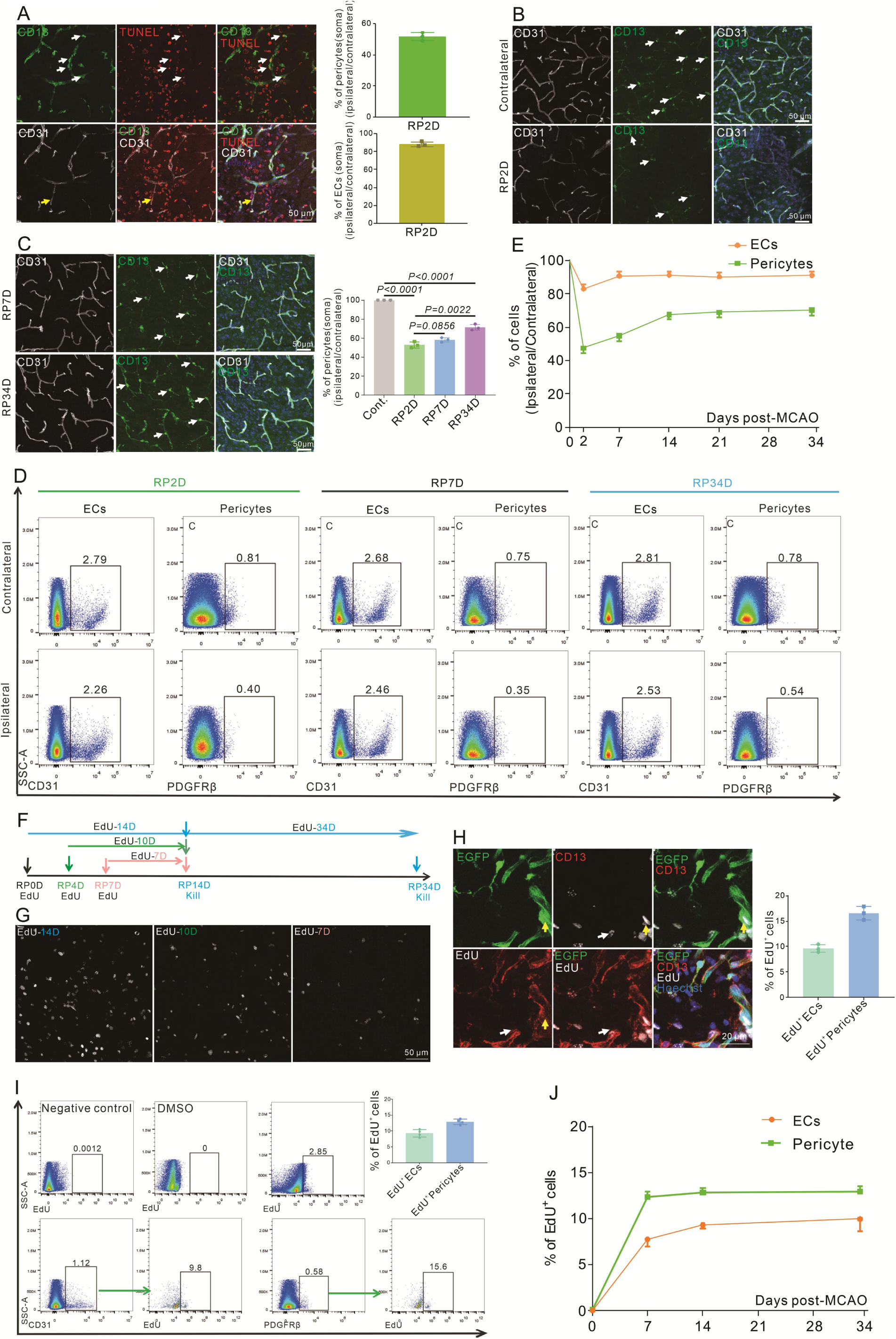
Pericytes die rapidly in the acute phase and replenish in the subacute and chronic phases after stroke. A.Immunoflurescence staining shows Tunel^+^ pericytes(white) and Tunel^+^ ECs(yellow) after MCAO at RP2D and quantitative the proportion of Tunel^+^ cells in pericytes and ECs(n=3, 20 slices/mouse). B.Immunoflurescence staining shows CD13^+^ soma after MCAO at RP2D(n=3,20 slices/mouse). C.Immunoflurescence staining shows CD13^+^ soma after MCAO at RP7D and RP34D, quantitative the ratio of CD13^+^ soma (n=3,20 slices/mouse). D.Flow cytometry analysis of the proportion of pericytes and ECs after MCAO at RP2D, RP7D and RP34D(n=6). E. Quantitative the proportion of pericytes and ECs at different reperfusion times after stroke(n=6). F.Schematic diagram displaying the time course for EdU injection and analysis time points. G.Maximum EdU signal in the ischemic area at different EdU injection times(n=3). H.Immunoflurescence staining shows EdU^+^ pericytes (white) and EdU^+^ ECs(yellow) after MCAO at RP34D and quantitative the proportion of EdU^+^ cells in pericytes and ECs(n=3,20 slices/mouse). I.Flow cytometry analysis of the proportion of EdU^+^ pericytes and EdU^+^ ECs after MCAO at RP14D (n=4). J.Flow cytometry analysis of the proportion of EdU^+^ pericytes and EdU^+^ ECs after MCAO at RP7D, RP14 and RP34D(n=4).

Considering these findings, we next aimed to assess why the number of pericytes increased in the chronic phase after stroke. EdU was injected into the mice at different times to identify proliferating pericytes and ECs (Figure 1F), and it was found that cell proliferation occurred mainly within the first three days after stroke (Figure 1G). Both pericytes and ECs underwent self-proliferation (Figure 1H), EdU^+^ ECs accounted for fewer than 10% of all ECs and EdU^+^ pericytes accounted for approximately 12.9% of all pericytes (Figure 1I). EdU^+^ pericytes and ECs ceased proliferating after RP7D (Figure 1J). Thus, 12.9% of the pericytes present at RP34D originated from self-proliferation, whereas the origin of the other 27.6% of new pericytes remained unknown.

### scRNA-seq was used to explore the fate of ECs after stroke

To explore the origin of the other 27.6% of new pericytes conversion from ECs after stroke, we used Cdh5CreERT2 ^36^(induced endothelial cells, iECs);Ai47 or Ai14 mice which were labeled ECs in homeostasis (Figure S1A). First, we confirmed that ECs were specifically labeled in the brain parenchyma of iECs;Ai47 mice. Without tamoxifen, ECs were not labeled with EGFP (Figure S1B-C). Immunofluorescence staining revealed the EGFP signal only represents ECs markers positive (CD31^+^, ERG^+^, GLUT1^+^, and VE-Cadherin^+^) cells in the brain parenchyma (Figure S1D-G, S1I). We also found that the EGFP signal does not represent pericyte markers (CD13^+^, PDGFRβ^+^, α-SMA^+^, and NG2^+^ cells) cells in the brain parenchyma (Figure S1H-J). Therefore, vascular ECs in the brain parenchyma were specifically labeled by EGFP from the homeostasis of iECs;Ai47 mice.

We isolated EGFP^+^ cells in the sham group of iECs;Ai47 mice and the MCAO groups at RP7D and RP34D. After sorting by 10X Genomics and quality control (Figure 2A, Figure S2A), we obtained 3568 cells from the sham group, 4147 cells from the MCAO group at RP7D and 5070 cells from the MCAO group at RP34D. We used different markers based on literature^37–39^ to identify the cell types (Figure 2B) and found 9 major cell types: arterial ECs, venous ECs (2 types), capillary ECs, capillary-venous ECs, smooth muscle cells (SMCs), pericytes, fibroblasts, microglia and ependymal cells (Figure 2C). Those cell types of highly expressed canonical markers: ECs expressed *Pecam1*, arterial ECs expressed *Gkn3*, venous ECs expressed *Slc38a5*, capillary ECs expressed *Rgcc*, capillary-venous ECs *Rar4*, SMCs expressed *Cnn1*, pericytes expressed *Pdgfrb*, fibroblasts expressed *Pdgfra*, microglia expressed *Iba1* and ependymal cells expressed *Dynlrb2* (Figure S2B). We also noted some unexpected cell types, which may come from the leptomeninges, choroid plexus, periventricular cells, peripheral blood in the ischemic area or contamination during sample processing. ECs accounted for nearly 90% of all cells for different groups (Figure S2C). Gene Ontology (GO) analysis of the top 50 differentially expressed genes (DEGs) (Figure 2D) from the 9 major groups showed the functional roles of the genes. GO analysis identified the biological processes, molecular functions, and cellular components associated with the DEGs in each group. We calculated the number of cells in the sham group and MCAO group at RP7D and RP34D, and the proportions of 3 cell types (pericytes, fibroblasts and microglia) markedly increased (Figure 2E, Figure S2C).

**Figure 2.**
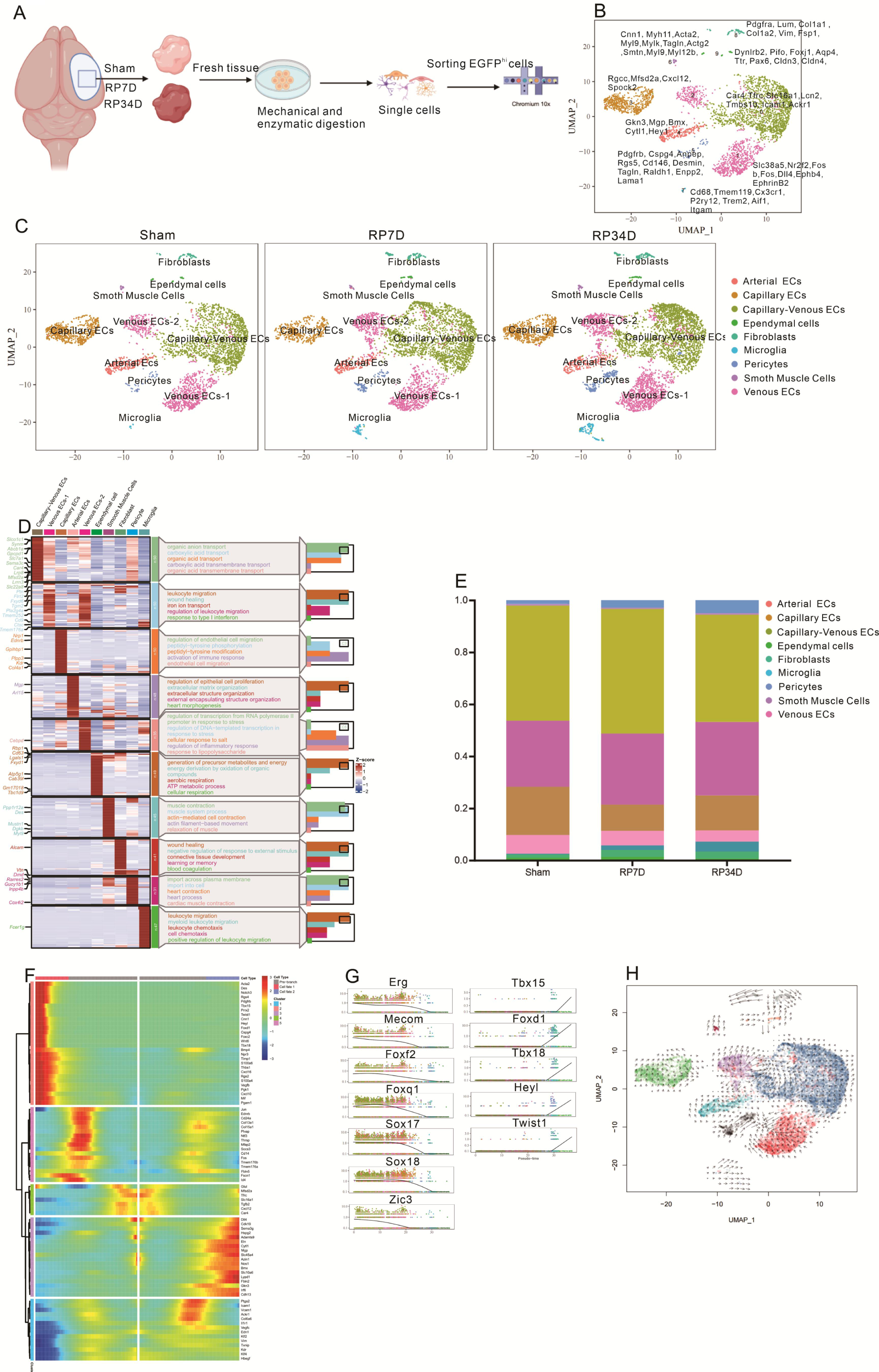
scRNA-seq is used to explore the fate of ECs after stroke. A.Samples were obtained from mouse ischemic brains at Sham, RP7D and RP34D. Single cells were processed using Chromium 10x 3′DEG chemistry. B.Uniform manifold approximation and projection (UMAP) embedding of all cells and marker genes. C.UMAP analysis of individual Sham, RP7D and RP34D cell transcriptomes showed 10 clusters. D.Heatmap displays an expression of the top 50 upregulated genes in each cluster. The scale bar represents the z-score of average gene expression (log). E.Relative proportion of major cell types in different reperfusion times. F.Differential genes expression variance over pseudotime of ECs differentiation trajectory branches. G.Differential transcription factors expression variance over pseudotime of ECs differentiation trajectory branches. H.Pseudotime trajectory of ECs differentiation trajectory branches. The arrows show the direction of pseudotime trajectories.

A heatmap was used to visualize the top DEGs in the pseudotime trajectory (Figure 2F) and found that they were related mainly to pericyte transcription factors^40^ (Tbx15, Foxc2, Twist1, Tbx18) and pericyte markers^40^ (Cspg4, Pdgfrb, Rgs4) in cell fate P1 and arterial ECs markers (Mgp, Gkn3) in cell fate P2. Among the DEGs, EC-related transcription factors^40^ (Erg, Mecom, Foxf2, Foxq1, Sox17, Sox18 and Zic3) were downregulated as differentiation progressed, and the expression of pericyte-related transcription factors (Tbx15, Foxd1, Tbx18, Heyl and Twist1), which are critical for pericyte development, increased (Figure 2G). Pseudotime trajectory analysis suggested that the developmental trajectory was from ECs to pericytes and arterial ECs (Figure 2H). Thus, the scRNA-seq results indicated that ECs might turn into pericyte-like cells after stroke.

### ECs can give rise to pericyte-like cells after stroke

To verify that ECs might give rise to pericyte-like cells after stroke, we performed lineage tracing in iECs;Ai47 mice subjected to stroke. The graph shows the time of given tamoxifen, MACO model and different time points for analysis (Figure S3A). Laser speckle analysis revealed that CBF was reduced by approximately 80% after MCAO, indicating our ischemic model success in mice (Figure S3B). Immunofluorescence revealed that there were existing EGFP^+^ cells, detached from blood vessels, owning to long processes with secondary branches and no expression EC markers (CD31, ERG, GLUT1 and VE-Cadherin), in the ischemic area at RP34D (Figure 3A, Figure S3C). Theoretically, EGFP^+^ cells only exist in ECs and express ECs markers in parenchyma at homeostasis. However, the above EGFP^+^ cells lost the characteristics of ECs (EGFP^+^ non-ECs), which indicated those cells had changed fate. Moreover, the EGFP^+^CD31^-^ cells expressed classic pericyte markers (CD13, PDGFRβ and NG2) (Figure 3A-C), which also indicated ECs might turn into pericyte-like cells after stroke.

**Figure 3.**
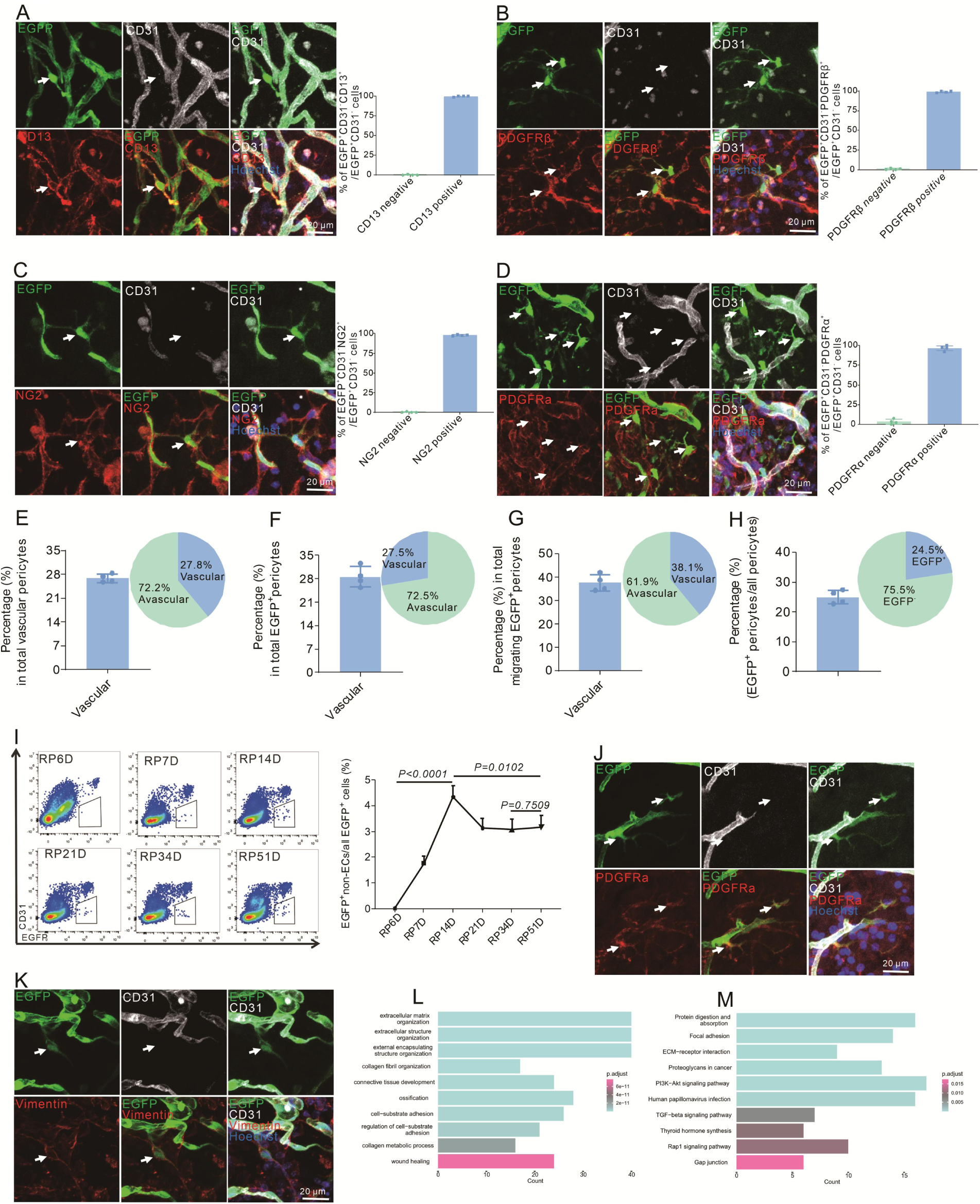
Endothelial-to-pericytic transition can replenish pericytes and undergo an intermediate fibroblast-like cell state after stroke. A.Immunoflurescence staining of CD31 and CD13 expression in Cdh5CreERT2;Ai47 mice with MCAO at RP34D and quantitative the proportion of CD13^+^ cells(n=5,20 slices/mouse). B.Immunoflurescence staining of CD31 and PDGFRβ expression in Cdh5CreERT2;Ai47 mice with MCAO at RP34D and quantitative the proportion of PDGFRβ^+^ cells(n=5,20 slices/mouse). C.Immunoflurescence staining of CD31 and NG2 expression in Cdh5CreERT2;Ai47 mice with MCAO at RP34D and quantitative the proportion of NG2^+^ cells(n=5,20 slices/mouse). D.Immunoflurescence staining of CD31 and PDGFRα expression in Cdh5CreERT2;Ai47 mice with MCAO at RP34D and quantitative the proportion of PDGFRα^+^ cells (n=5,20 slices/mouse). E.Quantitative the proportion of EGFP^+^ pericytes on blood vessel/pericytes on blood vessel(n=5,20 slices/mouse). F.Quantitative the proportion of EGFP^+^ pericytes on blood vessel/EGFP^+^ pericytes (n=5,20 slices/ mouse). G.Quantitative the proportion of EGFP^+^ pericytes on blood vessel/ migrating EGFP^+^ pericytes(n=5, 20 slices/mouse). H.Quantitative the proportion of EGFP^+^ pericytes/all pericytes(n=5, 20 slices/ mouse). I.Flow cytometry analysis of the proportion of EGFP^+^&CD31^-^ cells after MCAO at different times and quantitative the proportion of EGFP^+^&CD31^-^ cells(n=4). J.Immunoflurescence staining of CD31 and PDGFRα expression in Cdh5CreERT2;Ai47 mice with MCAO at RP8D(n=5,20 slices/mouse). K.Immunoflurescence staining of CD31 and vimentin expression in Cdh5CreERT2;Ai47 mice with MCAO at RP8D (n=5,20 slices/mouse). L.GO analysis of upregulated DEGs in EGFP^+^&CD31^-^ cells subgroup, compared with EGFP^+^&CD31^+^ cells from contralateral. M.KEGG pathway analysis of upregulated DEGs in EGFP^+^&CD31^-^ cells subgroup, compared with EGFP^+^&CD31^+^ cells from contralateral.

Cardiac pericytes upregulate the expression of fibrosis-related genes, exhibiting matrix-synthesizing and matrix-remodeling abilities after myocardial infarction. Some pericytes in the infarct region express fibroblast marker PDGFRɑ, which is beneficial for ECM in myocardial infarction^41^. Our model also found that EGFP^+^ non-ECs expressed fibroblast marker PDGFRɑ (Figure 3D). However, EGFP^+^ non-ECs did not express vimentin (Figure S3D), which indicates that the EGFP^+^ non-ECs were not fibroblasts, but pericyte-like cells that expressed certain proteins involved in matrix remodeling. Moreover, EGFP^+^ non-ECs did not express microglia markers (Iba1 and CD68) (Figure S3E), which indicated that EGFP^+^ non-ECs were not microglia.

The proportion of EGFP^+^ non-ECs located on vessels relative to total vascular pericytes was approximately 27.8% (Figure 3E), that relative to total EGFP^+^ non-ECs was approximately 27.5% (Figure 3F), that relative to migrating EGFP^+^ non-ECs was approximately 38.1% (Figure 3G), and that relative to total pericytes was 22.45% (Figure 3H). EGFP^+^ non-ECs began to appear at RP7D, and their proportion peaked at RP14D (Figure 3I). Furthermore, the EGFP^+^ non-ECs survived over 514 days after MCAO (Figure S3F). At RP8D, EGFP^+^ non-ECs expressed PDGFRα (Figure 3J) and vimentin (Figure 3K) but not CD13 (Figure S3G) or NG2 (Figure S3H). GO and KEGG analyses also revealed that EGFP^+^ non-ECs expressed genes involved in matrix synthesis, matrix remodeling and ossification at RP8D (Figure 3L-M). These results indicated that EGFP^+^ non-ECs presented fibroblast-like cells characteristics at RP8D and pericyte-like cells at RP34D. The majority of the EGFP^+^ non-ECs were located in the penumbra (Figure S3I), and they were not proliferation, as they were not labeled by EdU at RP34D (Figure S3J), which indicated that the EGFP^+^ non-ECs were converted from ECs via transdifferentiation.

In iECs;Ai47 mice, fewer than 1% of nucleated cells in bone marrow (BM) were labeled with EGFP, main myeloid cells^42^. Macrophages could invade the brain and develop pericytes in the early phase of vascular development in the CNS^43^. Therefore, we aimed to eliminate EGFP^+^ macrophages from the BM after stroke. We intracerebroventricularly injected newborn Cdh5CreERT2 P2 mice with AVV2/9-CAG-DIO-EGFP. EGFP^+^ non-ECs still expressed the pericyte marker CD13 at RP34D, with the percentage of EGFP^+^ non-ECs expressing CD13 approaching 100% at RP34D (Figure S3K). We also used other methods to specifically label ECs and to verify the conversion of ECs to pericytes. In Tie2Dre;Mfsd2aCrexER;Ai47 mice, ECs in the brain are specifically labeled, in which EC would give rise to CD13^+^ EGFP^+^ non-ECs at RP34D (Figure S3L). Similarly, AAV2/9-BI30-CAG-EGFP specifically infected ECs, and following injection, close to 100% of EGFP^+^ non-ECs still expressed CD13 at RP34D (Figure S3M).

To explore transcriptome characteristics of EGFP^+^ non-ECs at RP34D, we isolated EGFP^+^ non-ECs at RP34D for RNA-seq analysis (Figure S4A). Principal component analysis revealed that the cells clustered into ECs and EGFP^+^ non-ECs (Figure S4B). Analysis of a heatmap showing the top 100 DEGs between ECs and EGFP^+^ non-ECs (Figure S4C) revealed differences in their transcriptomes. The heatmap showed that EC-specific transcription factor genes^40^ (Figure S4D), EC-enriched transmembrane receptor genes^40^ (Figure S4E) and EC-enriched ligand genes^40^ (Figure S4F) were expressed in ECs but expressed or expressed at low levels in EGFP^+^ non-ECs. However, pericyte-specific transcription factor genes^40^ (Figure S4G), pericyte-enriched transmembrane receptor genes^40^ (Figure S4H) and pericyte-enriched ligand genes^40^ (Figure S4I) were not expressed or expressed at low levels in ECs but expressed in EGFP^+^ non-ECs. Therefore, ECs lost their EC-like transcriptome, and EGFP^+^ non-ECs acquired a pericyte-like transcriptome after stroke. A total of 379 and 220 genes were significantly upregulated and downregulated, respectively, in EGFP^+^ non-ECs compared with ECs at RP34D (Figure S4J). GO enrichment analysis of the upregulated DEGs in EGFP^+^ non-ECs was further performed, and according to the top 10 GO terms, the EGFP^+^ non-ECs presented greater aerobic capacity (Figure S4K) than did the ECs, which exhibited a stronger preference for anaerobic metabolism^44,45^. KEGG pathway analysis of the upregulated DEGs in EGFP^+^ non-ECs was further performed, and the upregulated DEGs were most significantly enriched in endothelial cell migration, epithelial cell migration, epithelial and tissue migration, which was consistent with the detachment of EGFP^+^ non-ECs from blood vessels (Figure S3C). These results confirmed that EGFP^+^ non-ECs exhibit pericyte-like characteristics and ECs give rise to pericyte-like cells after stroke. Therefore, we named EGFP^+^ non-ECs with E-pericytes, as they were derived from ECs.

### Depletion of E-pericytes impedes BBB repair

During development and in adulthood, pericytes maintain BBB function. Reduced pericyte coverage can increase Evans blue accumulation in mutant brain parenchyma^20,46^. Whether E-pericytes generation also contributes to preserving BBB function after stroke is unknown. To specifically deplete E-pericytes after stroke (Figure 4A), we first used AAV2/9-BI30-NG2 promotor-DIO-DeRed (DeRed) to label E-pericytes at RP34D (Figure 4B-C). Then we used AAV2/9-BI30-NG2 promoter-DIO-DTA (DTA) virus, which injected before MCAO (Figure 4A), to deplete E-pericytes (Figure 4D). We found that Evans blue leakage decreased with increasing reperfusion time and that there was no obvious Evans blue signal at RP34D (Figure 4E), which indicates self-repair of the BBB. Depletion of E-pericytes by DTA resulted in a significant increase in Evans blue leakage (Figure 4F) and trypan blue leakage (Figure 4G) at RP34D after stroke. Furthermore, brain atrophy was more pronounced following the depletion of E-pericytes by DTA at RP34D after stroke (Figure 4F).

**Figure 4.**
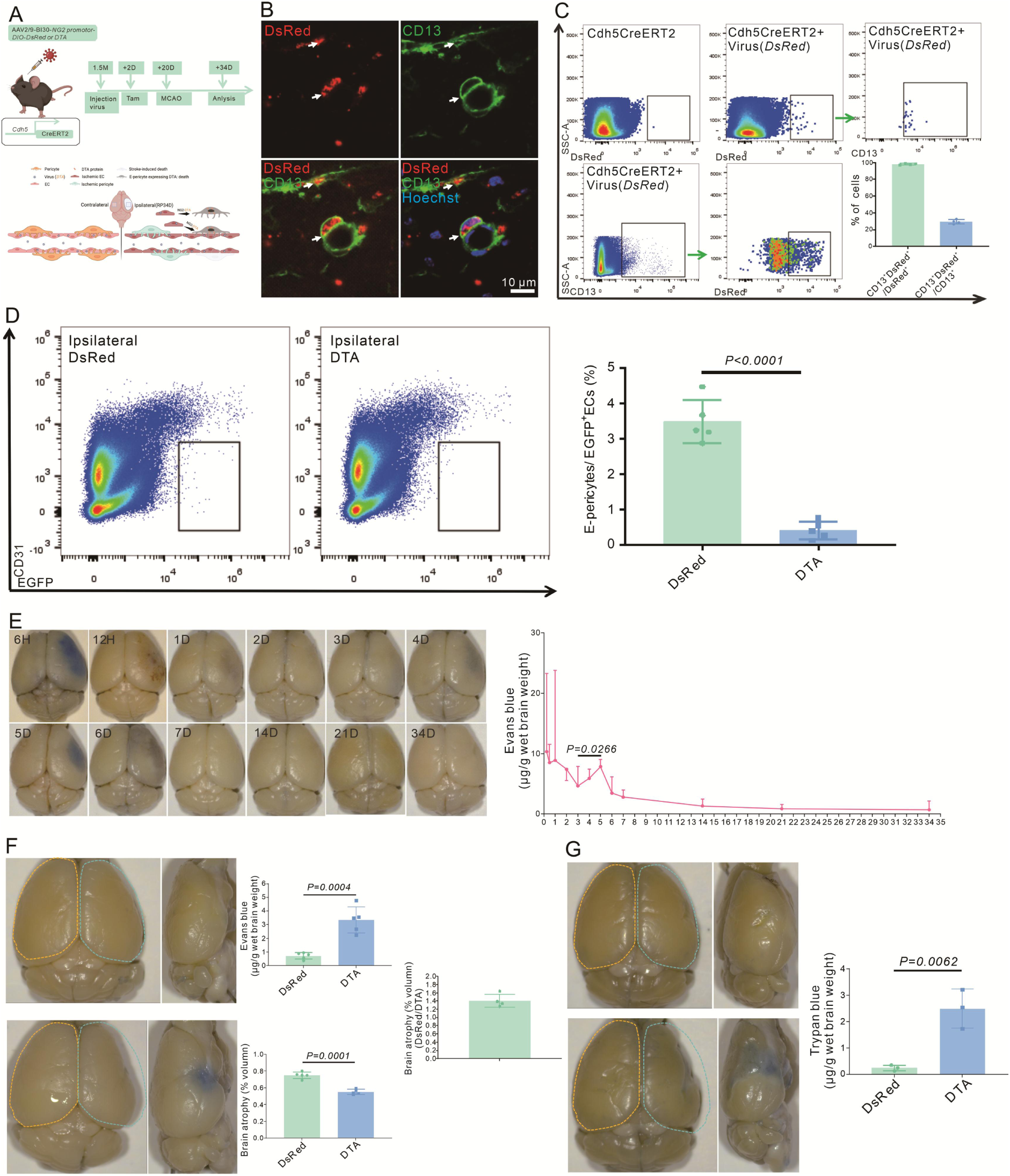
E-Perciytes deletion by AAV2/9-BI30-DIO-NG2-promotor-DTA virus aggravates BBB leakage after stroke. A.Schematic diagram displaying Cdh5CreERT2 injection with AAV2/9-BI30-DIO-NG2-promotor -DsRed or DTA virus to kill the cell and the time course for tamoxifen, MCAO and analysis time points. B.Immunoflurescence staining of CD13 expression in Cdh5CreERT2 injection with AAV2/9-BI30-DIO-NG2-promotor-DsRed A virus(n=3). C.Flow cytometry analysis the proportion of DsRed^+^&CD13^+^/ DsRed^+^ cells and quantitative the proportion (n=3). D.Flow cytometry analysis the proportion of E-Pericytes cells in Cdh5CreERT2 injection with AAV2/9-BI30-DIO-NG2-promotor-DTA virus and quantitative the proportion (n=5). E.Image showing the leakage of Evans blue in WT mice at different times after MCAO and quantitative the leakage of Evans blue(n=3). F.Image showing the leakage of Evans blue in Cdh5CreERT2 injection with AAV2/9-BI30-DIO-NG2-promotor-DTA virus at RP34D after MCAO, quantitative the leakage of Evans blue(n=5) and the brain atrophy volume (n=5). G.Image showing the leakage of trypan blue in Cdh5CreERT2 injection with AAV2/9-BI30-DIO-NG2-promotor-DTA virus at RP34D after MCAO and quantitative the leakage of trypan blue(n=3).

### Depletion of E-pericytes aggravates neurological deficits after stroke

Depletion of pericytes resulting in BBB dysfunction can impact normal brain function, damage neurons and impair cognitive and neurological functions^47,48^. E-pericyte generation after stroke may contribute to neuron survival and facilitate spontaneous behavioral recovery by restoration of BBB function.

Depletion of E-pericytes by DTA resulted in a decreased survival rate (Figure 5A), a decreased latency to fall from the rotarod in the rotarod test (Figure 5B), an increased frequency of falling from the pole (Figure 5C), an increased deviation from normal in the corner test (Figure 5D), and increased difficulty in removing adhesive tape from the forepaw in the adhesive removal test (Figure 5E). We detected a reduction in NeuN^+^ neurons after stroke when E-pericytes were depletion by DTA (Figure 5F). Furthermore, E-pericytes expressed higher levels of genes related to neuronal survival and growth, which are beneficial for neurons, compared to ECs (Figure 5G). The above results demonstrate that E-pericytes are involved in reducing neurological deficits and facilitating spontaneous behavioral recovery after stroke.

**Figure 5.**
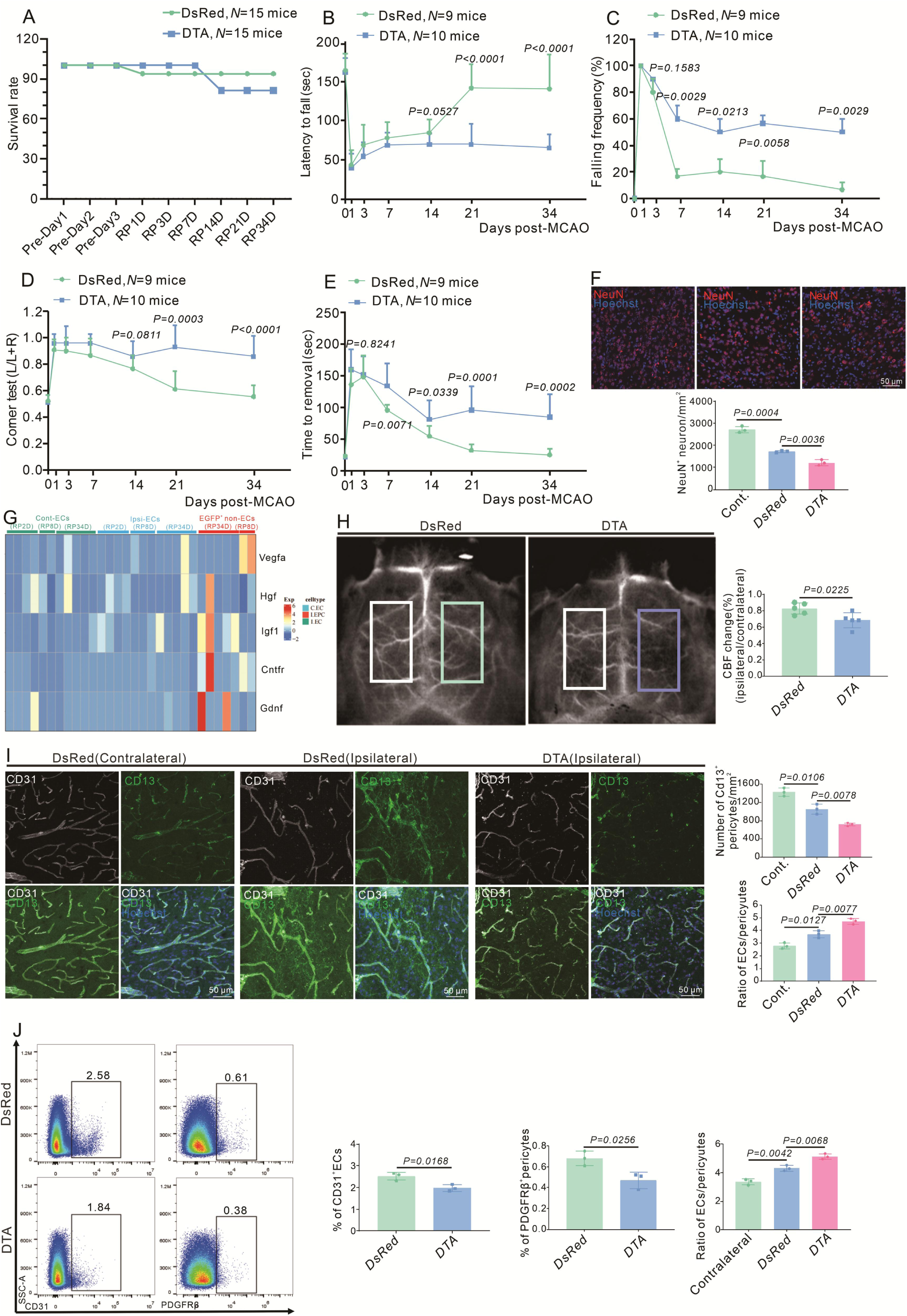
E-Pericytes deletion by AAV2/9-BI30-DIO-NG2-promotor-DTA virus exacerbates neurological deficit after stroke. A. Graph showing the survival rate of mice in Cdh5CreERT2 mice injection with AAV2/9-BI30-DIO-NG2-promotor-DsRed or DTA virus after MCAO at RP34D(n=15). B.Graph showing rotarod test in Cdh5CreERT2 mice injection with AAV2/9-BI30-DIO-NG2-promotor-DsRed or DTA virus after MCAO(n=9-10). C.Graph showing beam walking test in Cdh5CreERT2 mice injection with AAV2/9-BI30-DIO-NG2-promotor-DsRed or DTA virus after MCAO(n=9-10). D.Graph showing corner test in Cdh5CreERT2 mice injection with AAV2/9-BI30-DIO-NG2-promotor-DsRed or DTA virus after MCAO (n=9-10). E.Graph showing adhesive movement test in Cdh5CreERT2 mice injection with AAV2/9-BI30-DIO-NG2-promotor-DsRed or DTA virus after MCAO(n=9-10). F.Immunoflurescence staining of NeuN expression in Cdh5CreERT2 mice injection with AAV2/9-BI30-DIO-NG2-Long-DsRed or DTA virus after MCAO at RP34D and quantitative the number in the unit area(n=4,20 slices/mouse). G.The heat map showing promotion neuron survival and growth genes expression in all groups (n=5). H.Image showing the change of CBF in Cdh5CreERT2 mice injection with AAV2/9-BI30-DIO-NG2-promotor-DsRed or DTA virus after MCAO at RP34D and quantitative the change of CBF(n=5). I.Immunoflurescence staining of CD13 and CD31 expression in Cdh5CreERT2 mice injection with AAV2/9-BI30-DIO-NG2-promotor-DsRed or DTA virus after MCAO at RP34D and quantitative the percentage (n=3,20 slices/mouse). J.Flow cytometry analysis of the proportion of CD13^+^ cells and CD31^+^ cells in Cdh5CreERT2 mice injection with AAV2/9-BI30-DIO-NG2-promotor-DsRed or DTA virus after MCAO at RP34D and quantitative the percentage (n=3).

Additionally, when E-pericytes were depleted by DTA, there was a decrease in CBF at RP34D after stroke (Figure 5H). The number of CD31^+^ ECs and PDGFRβ^+^ pericytes decreased in immunofluorescence at RP34D after stroke when E-pericytes were depleted by DTA (Figure 5I-J), indicating decreased microvasculature. When E-pericytes were depleted, the number of CD13^+^ pericytes in the ischemic area was reduced by flow cytometry at RP34D after stroke, and the EC/pericyte ratio was increased (Figure 5I-J), suggesting that vascular integrity was disrupted. The above results demonstrated that E-pericytes are involved in keeping vascular functional integrity, relating to CBF and the EC/pericyte ratio.

### The TGFβ–TGFβR2 pathway impacts EndoMT and E-pericytes

Lineage tracing revealed that Procr^+^ ECs can give rise to pericytes during mammary gland development, and the transcriptome of Procr^+^ ECs exhibits the feature of EndoMT^24^. KEGG analysis revealed that the components of the TGF-β signaling pathway, the main driving force for EndoMT, were highly expressed in EGFP^+^ non-ECs at RP8D (Figure 3M). We suspected that EndoMT might be involved in E-pericytes formation. First, we showed that EndoMT occurs in the brain after stroke. The expression of EndoMT marker p-SMAD3^49^ increased in endothelial cells until RP8D and was decreased at RP34D after stroke (Figure 6A). At RP8D, we observed two adjacent EFGP^+^ cells on the same vessel, one CD31^+^ p-SMAD3^-^ cell with EC-like traits and a CD31^-^ p-SMAD3^+^ EGFP^+^ non-EC, implying that EndoMT had been occurring (Figure 6B). Moreover, the expression of EndoMT markers KLF4 and α-SMA^49^ was increased in ECs until RP8D and was decreased at RP34D after stroke (Figure S5A-B). The expression of α-SMA in ECs was increased, as shown by flow cytometry analysis of cytoplasmic gene expression (Figure S5C). EndoMT marker genes were also expressed in ECs on the ipsilateral side at RP2D (Figure S5D). All these findings indicated that ECs underwent EndoMT after stroke.

**Figure 6.**
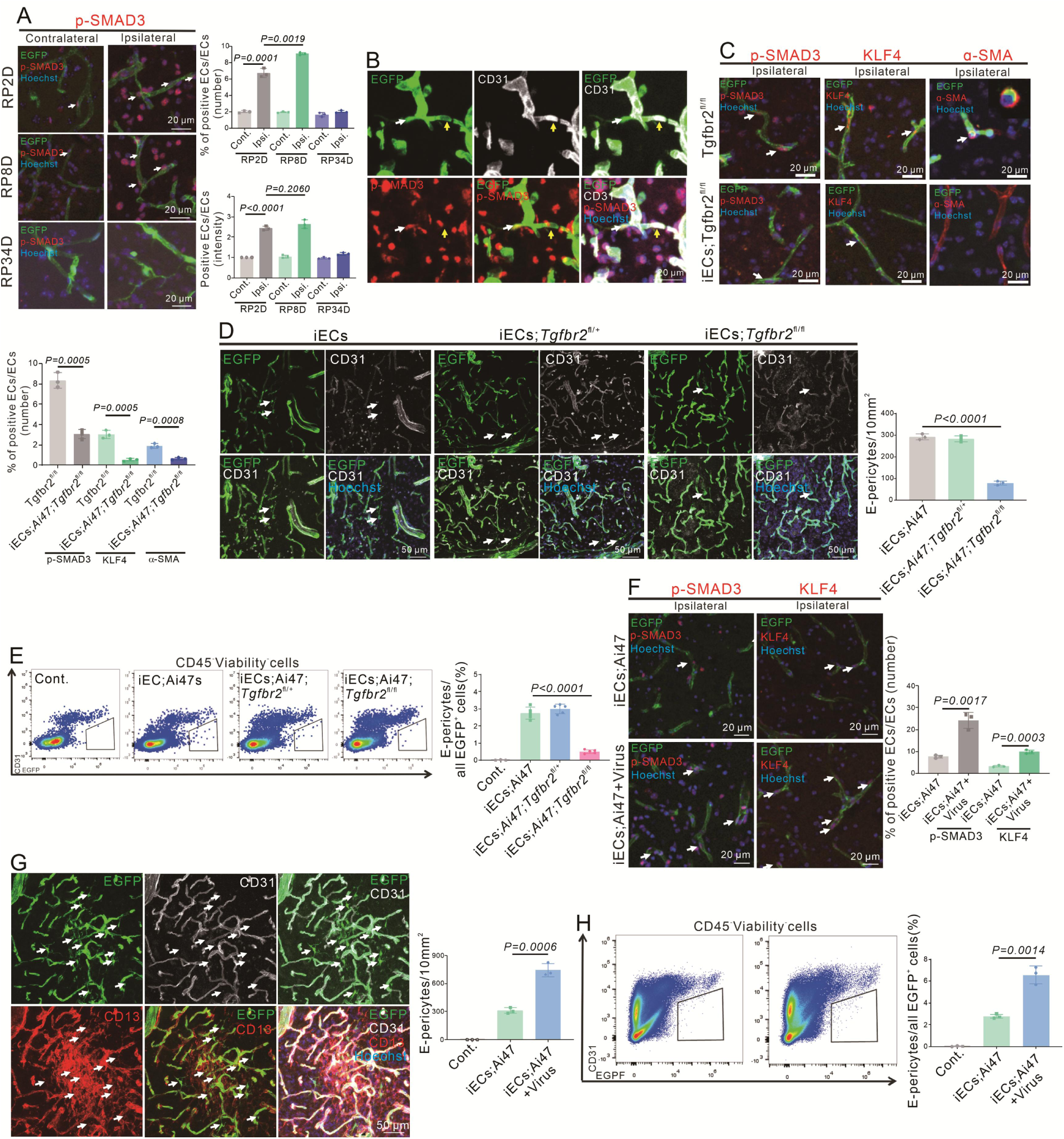
Endothelial-specific loss and reinforcement of Tgfbr2 gene expression affect EndoMT and E-Pericytes. A.Immunoflurescence staining of p-SMAD3 expression in Cdh5CreERT2;Ai47 mice at RP2D, RP8D and RP34D and quantitative the proportion and intensity(n=3,20 slices/mouse). B.Immunoflurescence staining of CD31 and p-SMAD3 expression in Cdh5CreERT2;Ai47 mice at RP8D (white:EGFP^+^&p-SMAD3^+^&CD31^-^; yellow:EGFP^+^&p-SMAD3^-^&CD31^+^; n=3). C.Immunoflurescence staining of EndoMT markers (p-SMAD3, KLF4 and α-SMA) expression in Cdh5CreERT2; Ai47;Tgfbr2^fl/fl^ mice at RP2D and quantitative the proportion and intensity(n=3,20 slices/mouse). D.Immunoflurescence staining of CD31 expression in Cdh5CreERT2; Ai47;Tgfbr2^fl/fl^ mice at RP34D and quantitative the number in the unit area (n=3,20 slices/mouse). E.Flow cytometry analysis the proportion of E-Pericytes in Cdh5CreERT2; Ai47;Tgfbr2^fl/fl^ mice at RP34D and quantitative the proportion(n=5). F.Immunoflurescence staining of EndoMT markers(p-SMAD3^+^ and KLF4^+^) expression in Cdh5CreERT2; Ai47 mice injection with AAV2/9-BI30-EF1α-DIO-Tgfbr2-3XFLAG-P2A-DsRed-WPREs at RP2D and quantitative the proportion of EndoMT markers(n=3,20 slices/mouse). G.Immunoflurescence staining of CD31 and CD13 expression in Cdh5CreERT2;Ai47 mice with AAV2/9-BI30-EF1α-DIO-Tgfbr2-3XFLAG-P2A-DsRed-WPREs at RP34D and quantitative the number in unit area(n=3,20 slices/mouse). H.Flow cytometry analysis of the proportion of E-Pericytes in Cdh5Cre ERT2; Ai47 mice with AAV2/9-BI30-EF1α-DIO-Tgfbr2-3XFLAG-P2A-DsRed-WPREs at RP34D and quantitative the proportion of E-Pericytes(n=3).

Additionally, flow cytometry analysis of cytoplasmic gene expression confirmed that 1.6% of CD31^+^ ECs expressed α-SMA at RP2 (Figure S5C), which was close to the proportion of E-pericytes relative to CD31^+^ ECs. Treatment with TGFβR2 inhibitors, L6293 and S1067, reduced the expression of EndoMT markers (Figure S5E), and endothelial cell-specific knockout of the *Tgfbr2* gene also decreased the expression of EndoMT markers (Figure 6C, Figure S5F). The TGFβR2 inhibitors diminished the number of E-pericytes (Figure S5G-H), and similarly, EC-specific knockout of the *Tgfbr2* gene diminished the proportion of E-pericytes (Figure 6D-E). Infection with *AAV2/9-BI30-EF1α-DIO-Tgfbr2-3XFLAG-P2A-DsRed-WPREs*[Virus*(Tgfbr2)*] resulted in EC-specific overexpression of the Tgfbr2 protein (Figure S5I), increased the expression of EndoMT markers in ECs (Figure 6F, Figure S5J) and promoted the generation of E-pericytes (Figure 6G-H). These findings suggested that stroke induces EndoMT and activation TGFβ-TGFβR2 signaling enhances EndoMT and E-pericytes, indicating that the endothelial-to-pericytic transition occurs via the TGFβ-EndoMT pathway.

### Infiltrating myeloid cells express TGFβ1 in the brain after stroke

We found that the TGFβ-TGFβR2 pathway is involved in the expression of EndoMT markers and the generation of E-pericytes after stroke. Therefore, we explored the source of TGFβ after stroke. The three main TGFβ isoforms in mammals are TGFβ1, TGFβ2, and TGFβ3, and TGFβ1 accounts for more than 90% of total TGFβ^50^. In young female mice subjected to distal middle cerebral artery occlusion (dMCAO), the TGFβ1 protein predominantly colocalized with CD68, activated microglia and macrophages, at 3 days poststroke^33^. RNAscope analysis revealed that *Tgfb1* mRNA was present predominantly (over 85%) in CD45^+^ immune cells at RP1D (Figure 7A) and RP2D (Figure S6A) but was present in only a fraction of Iba1^+^ microglia (∼15%) (Figure S6B). Moreover, *Tgfb1* mRNA was not expressed in CD13^+^ pericytes (Figure S6C). During the acute and subacute phases after stroke, immune cells that infiltrate the ischemic brain are primarily of the myeloid lineage^25^. Initially, flow cytometric analysis revealed that CD45^hi^ immune cells were predominantly of the myeloid lineage after stroke, with the majority being monocytes (Figure S6D-F). CD45^+^ cells were sorted 2 days after MCAO, and single cell sequencing analysis revealed that these cells were also primarily of the myeloid lineage (Figure 7B). Among myeloid cells, Monocyte-Derived Macrophages (MDM) were the main cells expressing *Tgfb1* mRNA (Figure 7C), with 96.3% of MDM expressing *Tgfb1* mRNA(Figure 7D). Immunofluorescence staining revealed almost no protein expression of TGFβ1 on the contralateral side (Figure 7E) but showed the presence of the TGFβ1 protein at RP2D; however, TGFβ1 protein expression was no longer observed at RP8D (Figure 7F). Furthermore, after anti-Ly6C/Ly6G antibodies were used to eliminate myeloid cells (Figure S6G-H), *Tgfb1* mRNA levels (Figure 7G) and Tgfb1 protein expression (Figure 7H) decreased. The above results demonstrated that myeloid cells could infiltrate the brain and release dominantly TGFβ1 after stroke.

**Figure 7.**
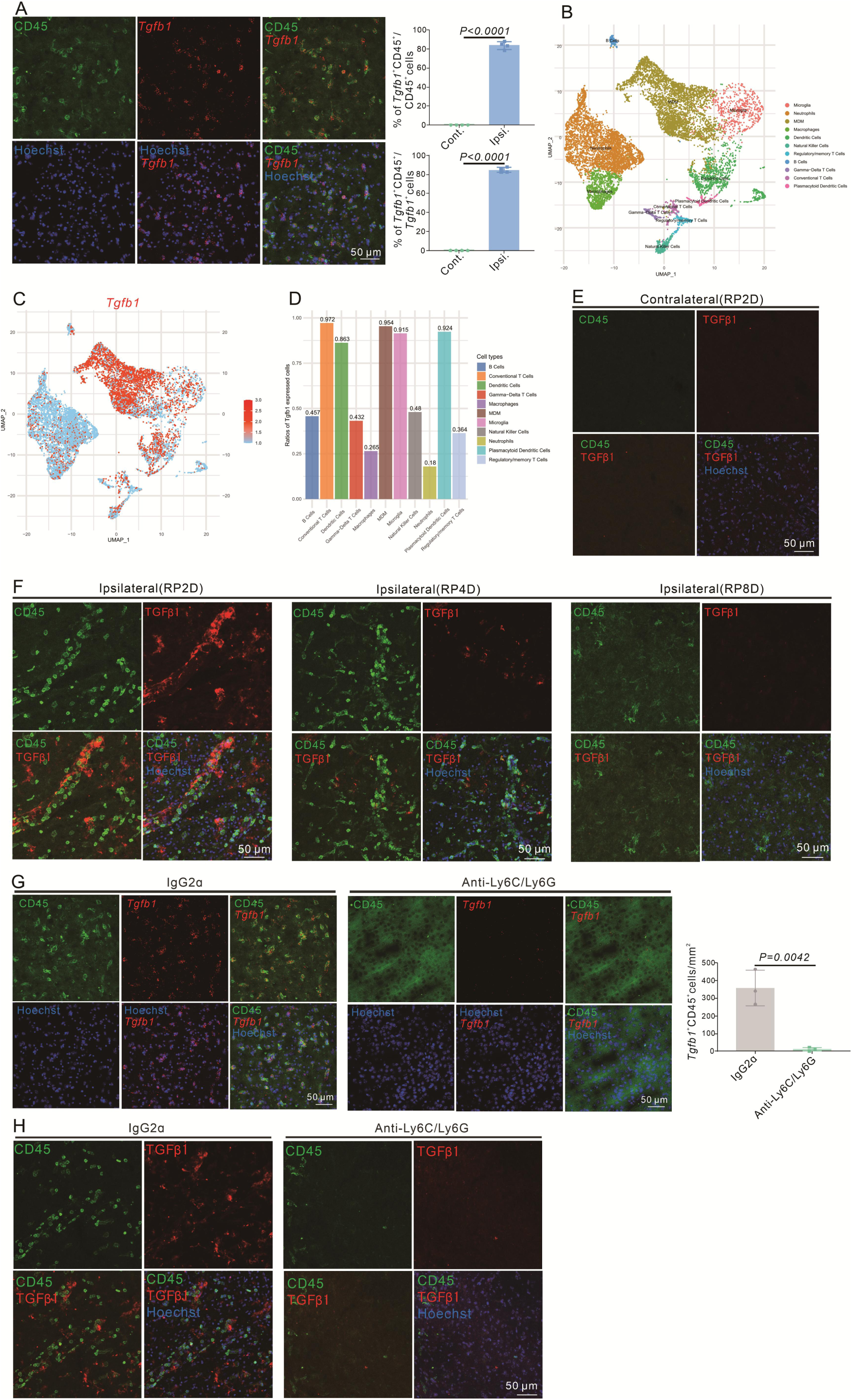
Infiltrating myeloid cells produce dominant TGFβ1 in mouse brain after stroke. A.Immunoflurescence staining of CD45 and *Tgfb1* gene expression in WT mice with MCAO at RP2D and quantitative the proportion of *Tgfb1*^+^ cells(n=4,10 slices/mouse). B.scRNA-seq of CD45^+^ cells isolated from the ipsilateral brain with MCAO:2H and RP1.5D. Dimensionality reduction and identification of clusters of transcriptionally similar cells were performed in an unsupervised manner(n=1), monocyte-derived macrophage (MDM). C.*Tgfb1* gene expression in dimensionality reduction and identification of clusters(n=1). D.The percentage of *Tgfb1* gene expression in different clusters(n=1). E.TGFβ1 protein expression in contralateral brain with MCAO at RP2D(n=3,20 slices/mouse). F.TGFβ1 protein expression in the ipsilateral brain with MCAO at RP2D, RP4D and RP8D (n=3,20 slices/mouse). G.Immunoflurescence staining of CD45 and *Tgfb1* gene expression in WT mice injection with anti-Ly6C/Ly6G after MCAO at RP2D and quantitative the proportion of CD45^+^&Tgfb1^+^ cells in the unit area(n=4,10 slices/mouse). H.Immunoflurescence staining of CD45 and Tgfb1 protein expression in WT mice injection with anti-Ly6C/Ly6G after MCAO at RP2D.

### Infiltrating myeloid cells drive the generation of E-pericytes, promote BBB recovery and neurological recovery after stroke

We found that myeloid cells could infiltrate the brain and release dominantly TGFβ1. Therefore, we explored whether myeloid cells could drive E-pericyte and promote BBB repair and neurological recovery after stroke.

First, the administration of anti-Ly6C/Ly6G antibodies to eliminate myeloid cells significantly reduced the number of CD45^hi^ immune cells in the brain at RP34D (Figure S7A-B), and it also decreased the generation of E-pericytes (Figure 8A-B). Furthermore, there was a significant increase in Evans blue extravasation, and brain atrophy was more pronounced at RP34D after stroke when infiltrating myeloid cells were eliminated (Figure 8C). These findings suggested that the inhibition of infiltrating myeloid cell-driven E-pericytes may have affected BBB function after stroke. CBF was decreased after infiltrating myeloid cells were eliminated at RP34D after stroke (Figure S7C). These findings indicated that the elimination of myeloid cells may have resulted in impaired restoration of CBF after stroke.

**Figure 8.**
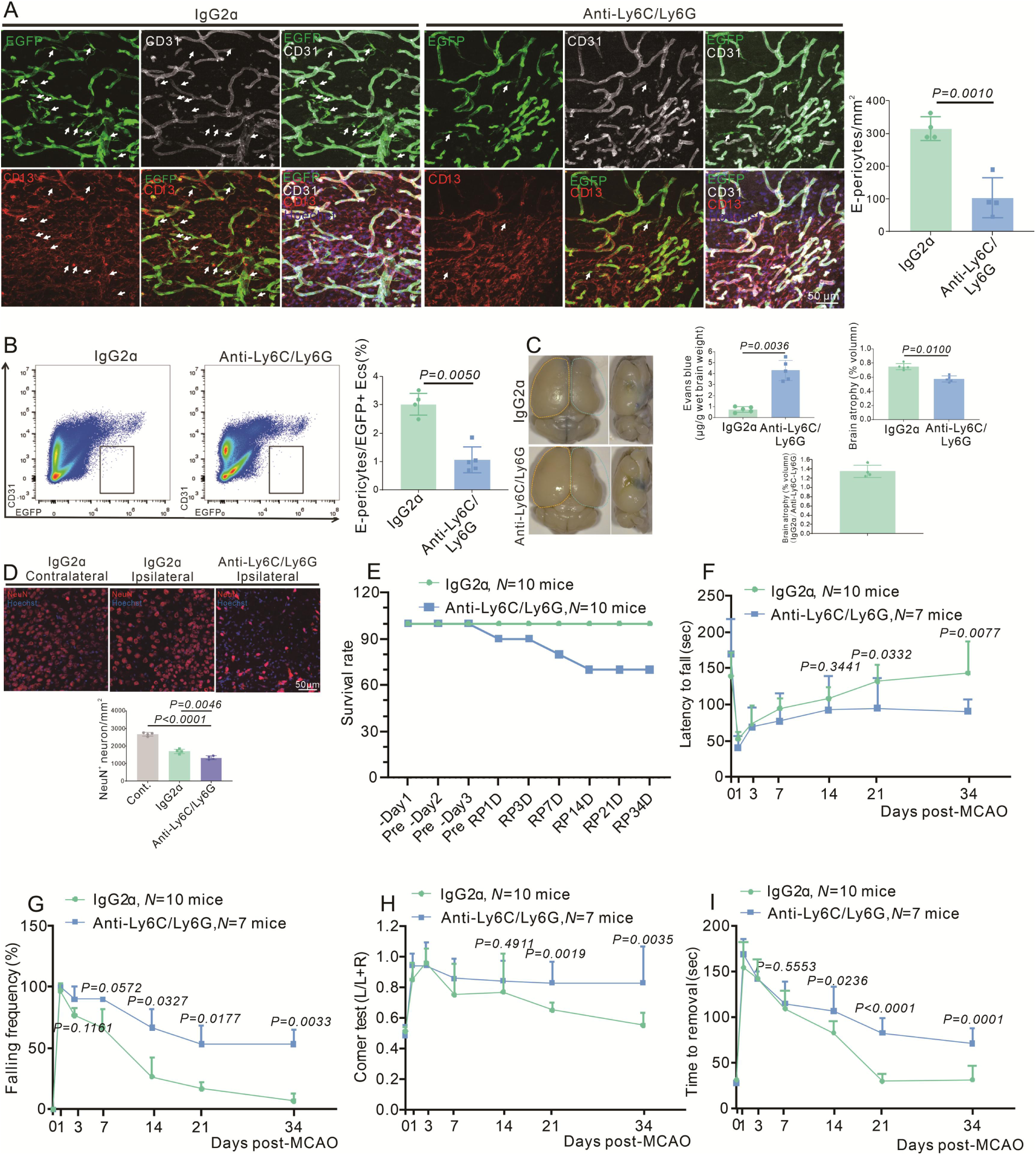
Infiltrating myeloid cells promote BBB renovation and neurological recovery after stroke. A.Immunoflurescence staining of CD13 and CD31 expression in WT mice injection with anti-Ly6C/Ly6G at RP34D after MCAO and quantitative the number in the unit area(n=4,20 slices/mouse). B.Flow cytometry analysis the proportion of EGFP^+^&CD31^+^ cells in WT mice injection with anti-Ly6C/Ly6G at RP34D after MCAO and quantitative the percentage(n=4). C.Image showing the leakage of Evans blue in WT mice injection with anti-Ly6C/Ly6G at RP34D after MCAO and quantitative the leakage of Evans blue and brain atrophy volume(n=5). D.Immunoflurescence staining of NeuN expression in WT mice injection with anti-Ly6C/Ly6G at RP34D after MCAO and quantitative the number in the unit area(n=4,20 slices/mouse). E.Graph showing the survival rate of mice in WT mice injection with anti-Ly6C/Ly6G at RP34D after MCAO(n=10). F.Graph showing rotarod test in WT mice injection with anti-Ly6C/Ly6G at RP34D after MCAO(n=7-10). G.Graph showing beam walking test in WT mice injection with anti-Ly6C/Ly6G at RP34D after MCAO(n=7-10). H.Graph showing corner test in WT mice injection with anti-Ly6C/Ly6G at RP34D after MCAO(n=7-10). I.Graph showing adhesive movement test in WT mice injection with anti-Ly6C/Ly6G at RP34D after MCAO(n=7-10).

We quantified the number of CD31^+^ ECs and found that it decreased when infiltrating myeloid cells were eliminated (Figure S7D), indicating decreased ECs. The number of CD13^+^ pericytes in the ischemic area was significantly reduced after ischemic stroke, and the EC/pericyte ratio was increased when infiltrating myeloid cells were eliminated (Figure S7D-E), which suggested that vascular integrity was disrupted. We also detected a reduction of NeuN^+^ neurons after ischemic stroke when infiltrating myeloid cells were eliminated (Figure 8D), indicating that fewer neurons survived. The elimination of infiltrating myeloid cells resulted in a low survival rate (Figure 8E). Moreover, the elimination of infiltrating myeloid cells decreased the latency to fall from the rotarod in the rotarod test (Figure 8F), increased the frequency of falling from the pole (Figure 8G), increased the deviation from normal in the corner test (Figure 8H), and impaired the removal of adhesive tape from the forepaw in the adhesive removal test (Figure 8I). The above results demonstrated that myeloid cells could infiltrate the brain and release TGFβ1, inducing the formation of E-pericytes after stroke. Thus, E-pericytes may be involved in the function of myeloid cells in promoting BBB restoration, reducing neurological deficits, and facilitating spontaneous behavioral recovery after stroke.

### EC-specific knockout of the *Tgfbr2* gene aggravates BBB leakage and neurological deficits after stroke

As we found that infiltrating myeloid cells drives the transdifferentiation of ECs into pericytes via the TGFb1-TGFβR2 pathway, we next aimed to explore whether blocking the TGFb1-TGFβR2 pathway in ECs also impacts BBB leakage and neurological deficits via E-pericytes. To this end, we increased the proportion of E-pericytes via EC-specific overexpression of the *Tgfbr2* gene or eliminated E-pericytes via EC-specific expression of the *DTA* gene. When Tgfbr2 was overexpressed to increase E-pericytes and DTA was expressed in transformed cells to deplete E-pericytes, we found that there was no significant change in the number of E-pericytes in the Tgfbr2 + DTA group compared with the DTA group (Figure 9A). Moreover, there was no difference in BBB leakage between the *DTA* expression group and the group in which *Tgfbr2* was overexpressed and *DTA* was expressed (Figure 9B). We found that Evans blue leakage increased at different time points after stroke in mice with EC-specific loss of the *Tgfbr2* gene, even up to RP118D after stroke (Figure 9C-D). Moreover, trypan blue leakage increased at RP34D with EC-specific loss of the *Tgfbr2* gene (Figure S8A). BBB leakage did not change significantly from RP7D to RP14D, while the proportion of E-pericytes increased. However, BBB leakage increased significantly after the elimination of E-pericytes (Figure 9D), which declares that E-pericytes promote the normal repair of BBB from RP7D to RP14D. Interestingly, no obvious dextran-rhodamine B (∼70 kDa) (Figure S8B) or Texas Red (∼71 kDa) leakage was detected (Figure S8C). The elimination of E-pericytes allowed Evans blue and trypan blue to cross the BBB. Thus, these findings indicate that E-pericytes contribute to BBB integrity. We detected a reduction in the number of NeuN^+^ neurons after stroke in mice with EC-specific loss of the *Tgfbr2* gene (Figure 9E), which led to much more severe neurological deficits. Specifically, EC-specific loss of the *Tgfbr2* gene in mice decreased the latency to fall from the rotarod (Figure 9F), increased the frequency of falling from the pole (Figure 9G), increased the deviation from normal in the corner test (Figure 9H), and impaired the removal of adhesive tape from the forepaw in the adhesive removal test (Figure 9I). The above results demonstrated that the TGFb1-TGFβR2 pathway induces the transdifferentiation of ECs into pericytes, contributing to BBB restoration and neurological recovery after stroke.

**Figure 9.**
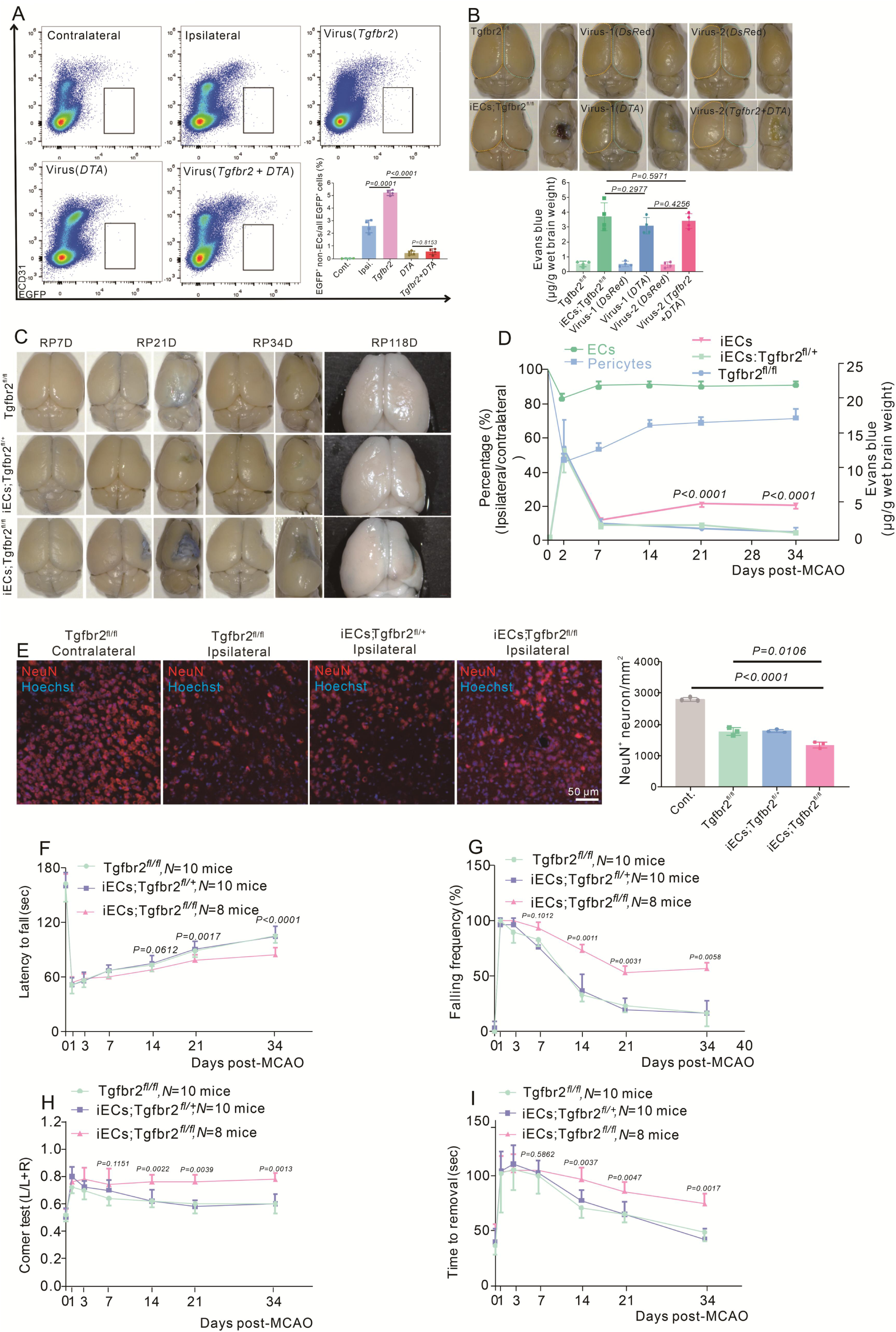
E-Pericytes deletion by specific endothelial cells knockout the *Tgfbr2* gene aggravates BBB leakage and neurologic deficit after stroke. A.Flow cytometry analysis of the proportion of E-Pericytes synchronous injection with AAV2/9-BI30-DIO-NG2-promotor DTA virus and AAV2/9-BI30-EF1α-DIO-Tgfbr2-3XFLAG-P2A-DsRed-WPREs virus after MCAO at 14D (n=4). B.Image showing the leakage of Evans blue in iECs;Tgfbr2^fl/fl^ mice at different times after MCAO and quantitative the leakage of Evans blue(n=3). C.Graph showing the leakage of Evans blue in iECs;Tgfbr2^fl/fl^ mice at different times after MCAO and quantitative the leakage of Trypan blue(n=3). D.Graph showing the percentage of pericyte and ECs, the leakage of Evans blue in iECs;Tgfbr2^fl/fl^ mice at different times after MCAO(n=3-6). E.Immunoflurescence staining of NeuN expression in iECs;Tgfbr2^fl/fl^ mice at RP34D after MCAO and quantitative the number of neurons in the unit area(n=3,20 slices/mouse). F.Graph showing the rotarod test in iECs;Tgfbr2^fl/fl^ mice after MCAO at RP34D(n=8-10). G.Graph showing the beam walking test in iECs;Tgfbr2^fl/fl^ mice after MCAO at RP34D(n=8-10). H.Graph showing the corner test in iECs;Tgfbr2^fl/fl^ mice after MCAO at RP34D(n=8-10). I.Graph showing the adhesive movement test in iECs; Tgfbr2^fl/fl^ mice after MCAO at RP34D (n=8-10).

### EC-specific overexpression of the *Tgfbr2* gene decreases BBB leakage and facilitates neurological recovery after stroke

The Elimination of E-pericytes by DTA exacerbated BBB leakage, decreased CBF, and impeded neurological and spontaneous behavioral recovery. Thus, increasing the number of E-pericytes may accelerate BBB repair, increase CBF, alleviate neurological deficits and promote spontaneous behavioral recovery after stroke. We previously found that EC-specific overexpression of *Tgfbr2* by *AAV2/9-BI30-EF1α-DIO-Tgfbr2-3XFLAG-P2A-DsRed-WPREs* promoted the generation of E-pericytes (Figure 6G-H). Additionally, EC-specific overexpression of the Tgfbr2 protein significantly increased CBF recovery (Figure S9A-B) and effective perfusion in the ischemic brain region at RP34D (Figure S9C-D). Thus, EC-specific overexpression of the Tgfbr2 protein promoted CBF restoration after stroke, through the generation of E-pericytes.

We quantified the number of CD31^+^ ECs and PDGFRβ^+^ pericytes and found that EC-specific overexpression of the Tgfbr2 protein by a virus (*Tgfbr2*) increased the number of these cells (Figure S9E) and the EC/pericyte ratio was close to normal (Figure S9E). EC-specific overexpression of the Tgfbr2 protein also decreased Evans blue leakage into the ischemic area (Figure 10A), although no obvious Evans blue signal was detected under bright field microscopy. Moreover, brain atrophy was reduced (Figure 10A).

**Figure 10.**
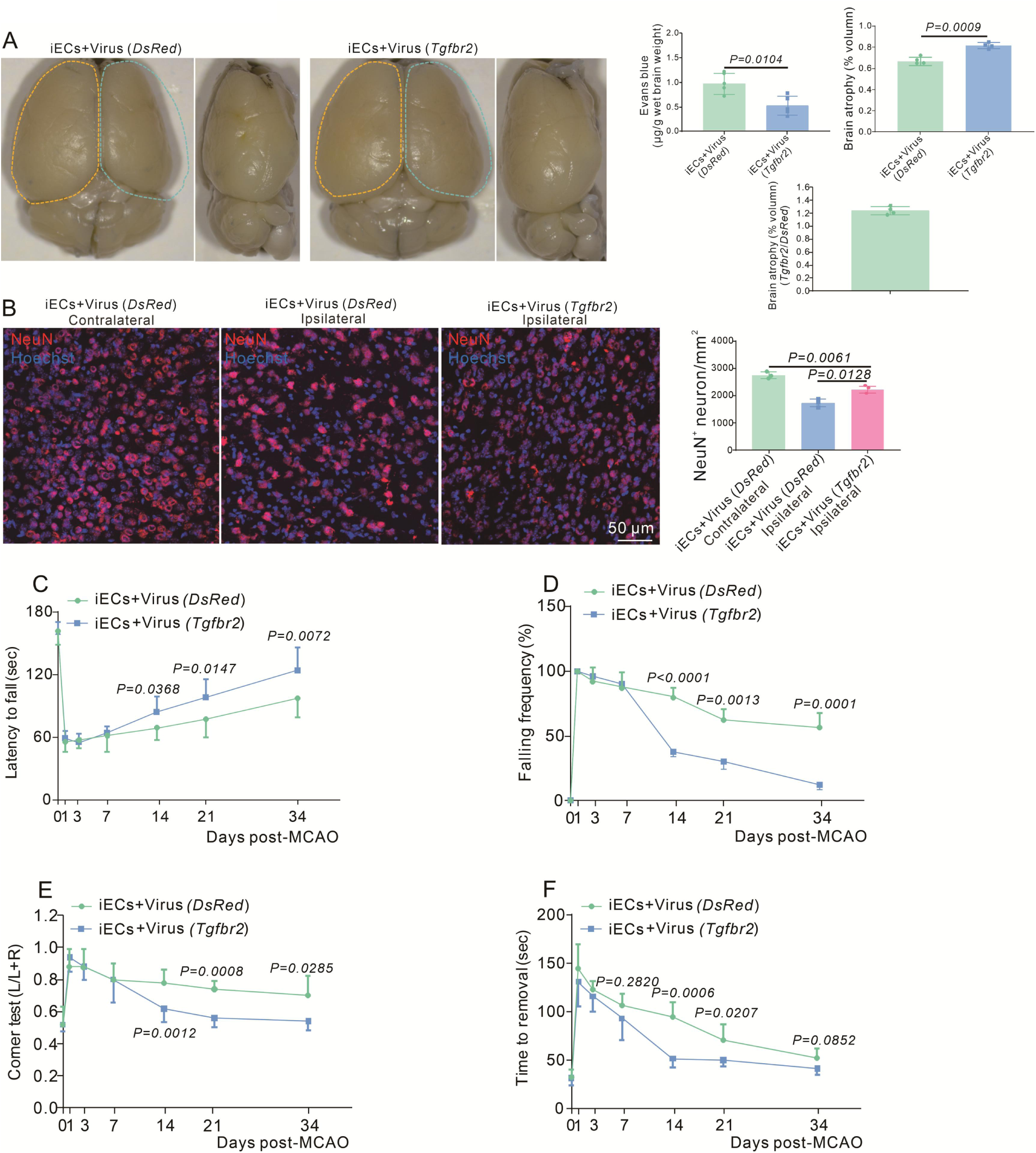
EC-specific overexpression of the *Tgfbr2* gene reinforces BBB function and neurological recovery after stroke. A. Image showing the leakage of Evans blue in Cdh5CreERT2 injection with AAV2/9-BI30-EF1α-DIO-Tgfbr2-3XFLAG-P2A-DsRed-WPREs virus at RP34D after MCAO and quantitative the leakage of Evans blue(n=5). B.Immunoflurescence staining of NeuN expression in Cdh5CreERT2 mice injection with AAV2/9-BI30-EF1α-DIO-Tgfbr2-3XFLAG-P2A-DsRed-WPREs virus after MCAO at RP34D and quantitative the number in the unit area(n=3,20 slices/mouse). C.Graph showing rotarod test in Cdh5CreERT2 mice injection with AAV2/9-BI30-EF1α-DIO-Tgfbr2-3XFLAG-P2A-DsRed-WPREs virus after MCAO(n=8-10). D.Graph showing beam walking test in Cdh5CreERT2 mice injection with AAV2/9-BI30-EF1α-DIO-Tgfbr2-3XFLAG-P2A-DsRedWPREs virus after MCAO(n=8-10). E.Graph showing corner test in Cdh5CreERT2 mice injection with AAV2/9-BI30-EF1α-DIO-Tgfbr2-3XFLAG-P2A-DsRed-WPREs virus after MCAO(n=8-10). F.Graph showing adhesive movement test in Cdh5CreERT2 mice injection with AAV2/9-BI30-EF1α-DIO-Tgfbr2-3XFLAG -P2A-DsRed-WPREs virus after MCAO(n=8-10).

Regarding neuronal changes, we also detected an increase of NeuN^+^ neurons after ischemic stroke in EC-specific overexpression of the Tgfbr2 protein group, indicating that more neurons survived (Figure 10B). Neurological deficits were assessed by evaluating locomotion after reperfusion at different times. EC-specific overexpression of the Tgfbr2 protein increased the latency to fall from the rotarod (Figure 10C) and decreased the frequency of falling from the pole in the rotating beam test (Figure 10D). Furthermore, EC-specific overexpression of the Tgfbr2 protein decreased the deviation from normal in the corner test (Figure 10E) and improved the ability of mice to remove adhesive tape from the forepaws (Figure 10F). These results suggested that EC-specific overexpression of the Tgfbr2 protein by a virus (*Tgfbr2*) decreases Evans blue leakage, promotes CBF recovery, alleviates neurological deficits and facilitates spontaneous behavioral recovery after stroke by increasing the number of E-pericytes.

## Discussion

This study demonstrated that ECs transdifferentiated into pericytes after stroke, accelerating BBB functional recovery, increasing CBF, and facilitating neurological behavioral recovery. Infiltrating myeloid cells were found to drive E-pericytes generation and restore BBB function and neurological functional recovery.

In other tissues, myeloid cells have been reported in tissue repair. Monocytes regulate hypodermal adipocytes and associated leptin-mediated revascularization of wounds post-infection^51^. Macrophages actively secrete metabolites during regeneration to establish immune–muscle stem cells (MuSCs) crosstalk in a GSDMD-dependent manner to promote tissue repair^52^. In our study, cerebral ischemic injury and repair also involve both the cerebral cells and the immune system, which play a critical role in determining stroke outcomes. Studies have shown that *CCR2* knockout and pharmacological inhibition of CCR2 attenuated monocyte-derived macrophage (MDM)-induced acute brain injury^30^. However, MDMs recruited to the injured brain early after ischemic stroke contribute to long-term spontaneous functional recovery during the chronic stage^30,32^. The ability of MDMs to promote functional recovery because macrophages are polarized toward the anti-inflammatory phenotype and promote angiogenesis during the recovery stage^30,32^. However, the precise role of myeloid cells in angiogenesis and spontaneous behavioral recovery after stroke remains to be elucidated. Our experimental results demonstrated that inhibiting the infiltration of myeloid cells into the brain could reduce the number of CD31^+^ ECs, implying the decrease of angiogenesis, and CBF. Furthermore, the suppression of myeloid cell-derived TGFβ1 in the brain restrains endothelial-to-pericytic transition and decreases the pool of pericytes. Pericytes play a vital role in angiogenesis and maintaining the stability of newly formed blood vessels. Therefore, the reduced number of pericytes caused by the inhibition of myeloid cells may ultimately impede angiogenesis and CBF. Our results may reveal that myeloid cells promote angiogenesis by driving the endothelial-to-pericytic transition.

During the acute phase of stroke, ischemia and hypoxia lead to rapid death of pericytes, which increases BBB leakage^20,53^. Although the decrease of pericytes during the acute phase of stroke has been reported, there is almost no research on the pericyte pool during the chronic phase. Our study found that during the chronic phase of stroke, the pericyte pool gradually increased, including both the self-proliferation of pericytes and the conversion of ECs to pericytes. E-pericytes were crucial for the function of vascular vessels during brain self-repair. When we depleted E-pericytes, it led to the loss of vascular vessels’ self-recovery function, including BBB and CBF, and hindered neurological recovery. Increasing the number of E-pericytes reduced Evans blue leakage, augmented CBF and promoted neurological recovery. This suggests the importance of E-pericytes for blood vessel self-recovery.

Unlike the liver and skin owing to powerful self-healing ability, stroke can lead to severe disability due to the limited capacity of cerebral cells to regenerate or replace dead cells, especially neurons. Exploring and confirming therapeutic theory for neurons and other cell protection is necessary for stroke. Synapse-level reconstruction of neural circuits and remyelination are involved in the recoverable process via various neurotrophic factors and cell transplants, which also are confronted with rigorous challenges to maintain long-term survival because of lack of effective blood flow^54,55^. Our results illustrated a new pathway for brain-self repair after stroke in the chronic stage. We found that ECs can give rise to pericytes to replenish the pool, accelerating the restoration of BBB function. Not only that, but E-pericytes may also increase angiogenesis, regulating no-reflow vessels to restore blood flow, thereby increasing the CBF of the whole ischemic brain. All the above improvements are beneficial in promoting neuron survival and functional recovery after stroke.

There is no doubt that the brain also has a limited ability to repair itself, which is not as strong as the liver and skin, but if you can amplify the brain’s ability to repair itself, it will be a huge progress for the therapy of stroke. The role of TGFβ family members in angiogenesis is underscored by the observations that nearly all mice with knockout of specific TGFβ family members die during midgestation due to yolk sac angiogenesis defects and that mutations in the genes encoding TGFβ pathway components are linked to an increasing number of human pathologies involving vascular dysfunction^56,57^. Our results revealed that increasing TGFβR2 expression could increase the number of E-pericytes, reduce BBB leakage, increase CBF and accelerate neurological functional recovery. Thus, the TGFβ1-TGFβR2 pathway is a potential therapeutic agent for promoting brain-self repair after stroke. At the same time, ECs may be transformed into pericytes in other tissues after ischemia, suggesting that activation of the TGFβ1-TGFβR2 pathway may also be involved in the repair of other tissues.

In summary, we presented transdifferentiation from ECs to pericytes after stroke, spanning from the processes of conversion to the functions of restoring BBB function and neurological functional recovery. We also verified that E-pericytes were driven by infiltrating myeloid cells, which released TGFβ1 to induce ECs subjected to EndoMT. The discovery of E-pericytes in fueling the destructive pericyte pool to remedy the functions of the lost pericytes after stroke indicates the capacity of brain self-repair from cell transformation.

### Limitations of the study

Our research revealed that after stroke, ECs can be converted into E-Pericytes. The TGFβ-TGFβR2 pathway had been shown to induce EndoMT in ECs, leading to the formation of fibroblast-like cells at RP8D. This phenomenon has been confirmed in other disease models. However, the conversion of fibroblast-like cells into pericytes has been rarely reported. We were not exploring how fibroblast-like cells convert into pericytes, which key transcription factors were involved in this process, and whether these transcription factors can be manipulated to regulate E-Pericytes more specifically, thereby promoting recovery after stroke. Some E-Pericytes were found to migrate from blood vessels, while others adhered with blood vessels. Based on the normal positioning of pericytes, E-Pericytes located on blood vessels are expected to function more effectively. Therefore, how to promote the return of migrating E-pericytes to blood vessels remains an unsolved problem.

---

**Figure S1.**
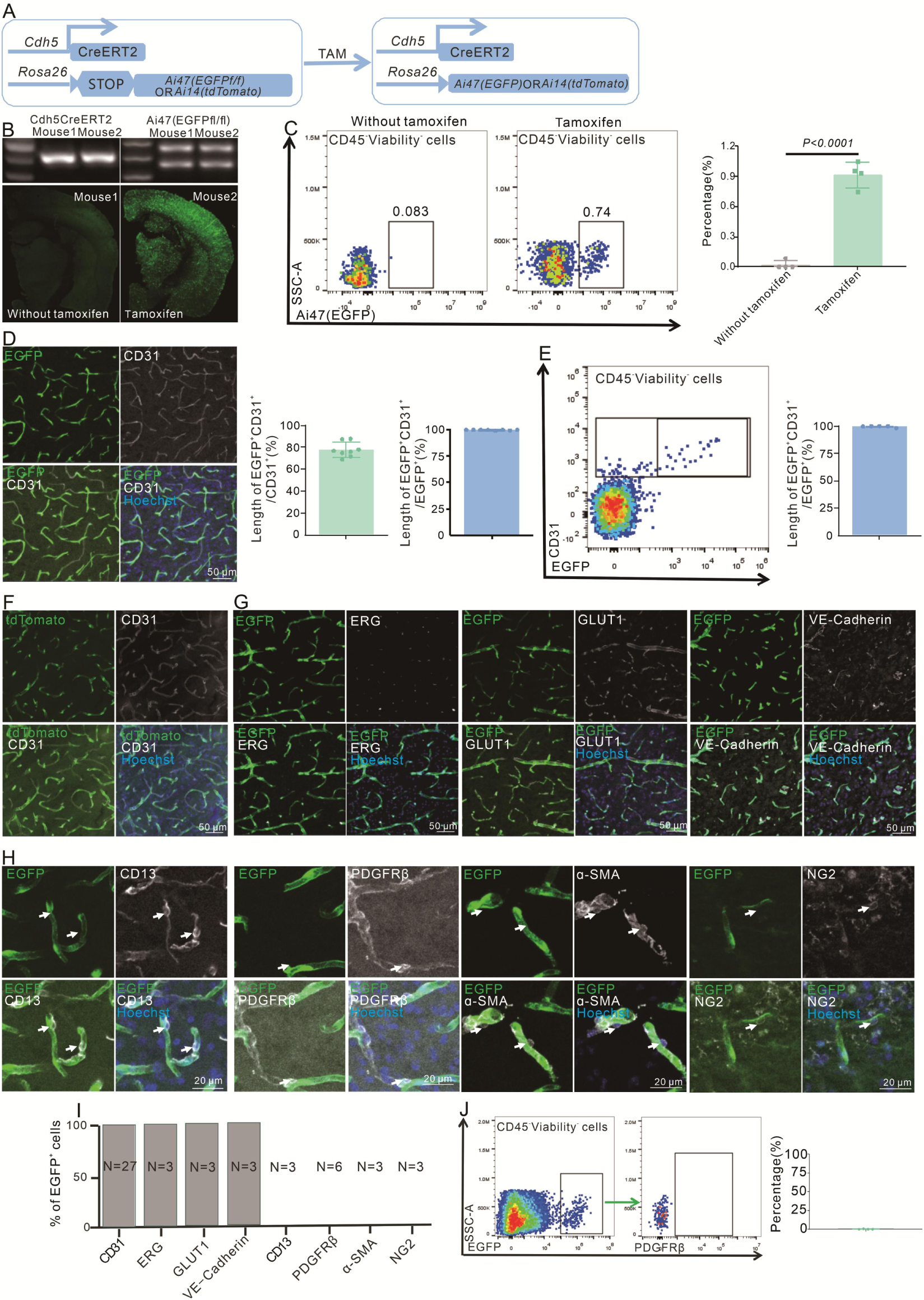
Cdh5CreERT2 mice specifically recombine parenchymal endothelial cells, related to. Figure 2. A.Schematic diagram displaying Cdh5CreERT2 mice label ECs. B.Immunoflurescence staining show with or without Tamoxifen in Cdh5Cre-ERT2; Ai47 mice(n=3,10 slices/mouse). C.Flow cytometry analyzes the proportion of EGFP^+^ cells and quantitatively the proportion of EGFP^+^ cells(n=4). D.Immunoflurescence staining of CD31 expression in Cdh5CreERT2;Ai47 mice and quantitative the proportion of EGFP^+^ &CD31^+^/ CD31^+^ or EGFP^+^ blood vessel(n=8,10 slices/mouse). E.Flow cytometry analysis of the proportion of EGFP^+^ &CD31^+^ /EGFP^+^ cells and quantitative the proportion(n=5). F.Immunoflurescence staining of CD31 expression in Cdh5CreERT2; Ai14 mice(n=3,20 slices/mouse). G. Immunofluorescence staining ECs markers (ERG, GLUT1 and VE-Cadherin) expression in Cdh5CreERT2; Ai47 mice (n=3-27, 10 slices/mouse). H.Immunoflurescence staining pericyte markers(CD13, PDGFRβ, α-SMA and NG2) expression in Cdh5CreERT2; Ai47 mice(n=3-6, 20 slices/mouse). I.Quantitative the proportion of F, G and H. J.Flow cytometry analysis of the proportion of PDGFRβ^+^ cells in EGFP^+^ cells and quantitative the proportion (n=4).

**Figure S2.**
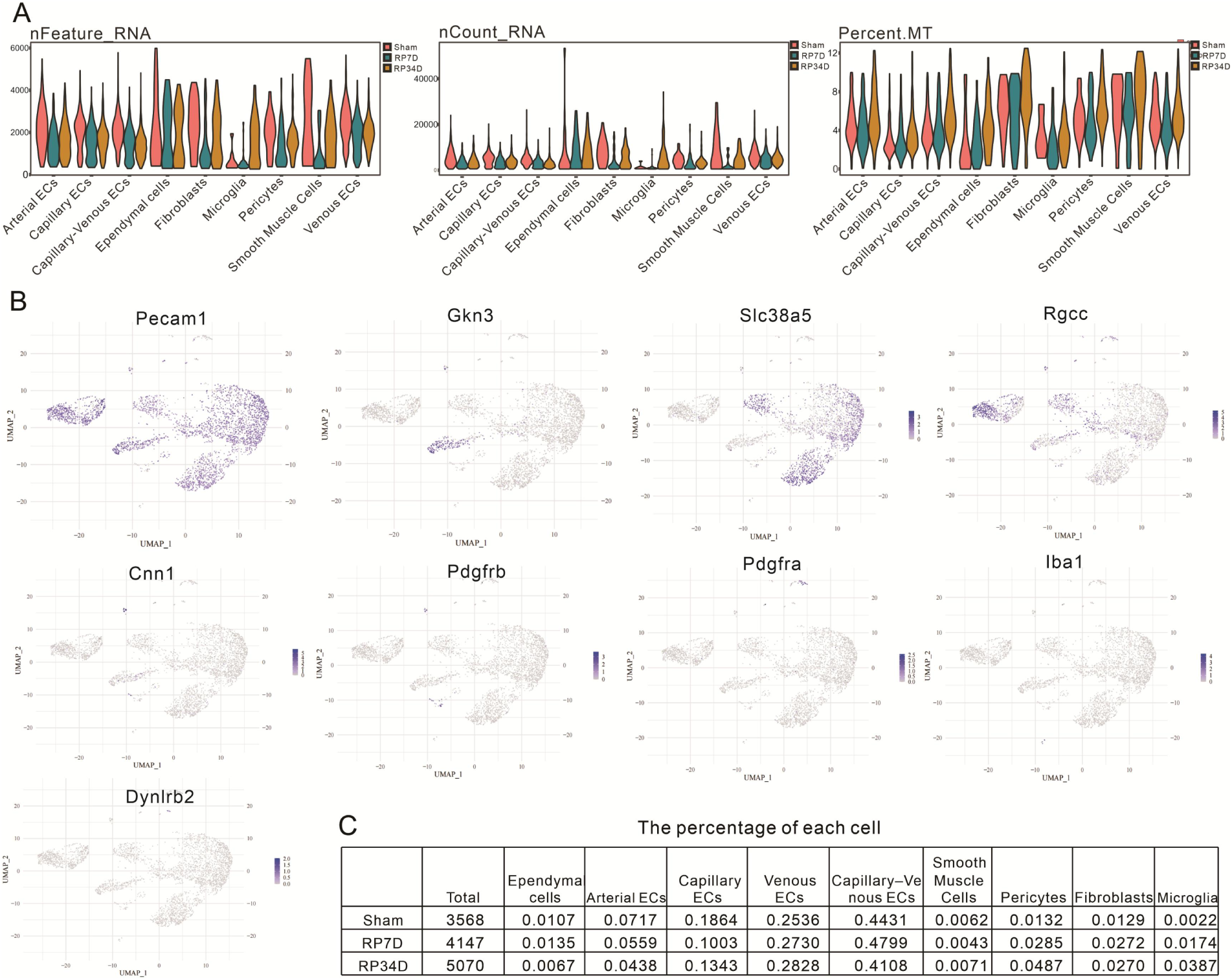
EGFP^+^ cells transcriptomic dataset quality. A.Violin plots showing the distribution of the number of total UMI counts per cell (nCount), genes detected per cell (nFeature), and percentage of mitochondrial genes (percent.mt) per identified cell type. B.UMAP plots depicting expression of individual marker genes for ECs (Pecam1), Arterial ECs (Gkn3), Venous ECs (Slc38a5), Capillary ECs (Rgcc), SMCs (Cnn1), Pericytes (Pdgfrb), Fibroblasts (Pdgfra), Microglia (Iba1) and Ependymal cells (Dynlrb2). Scale bars represent the log of normalized gene expression. C.Statistical table showing the percentage and number for each cell type.

**Figure S3.**
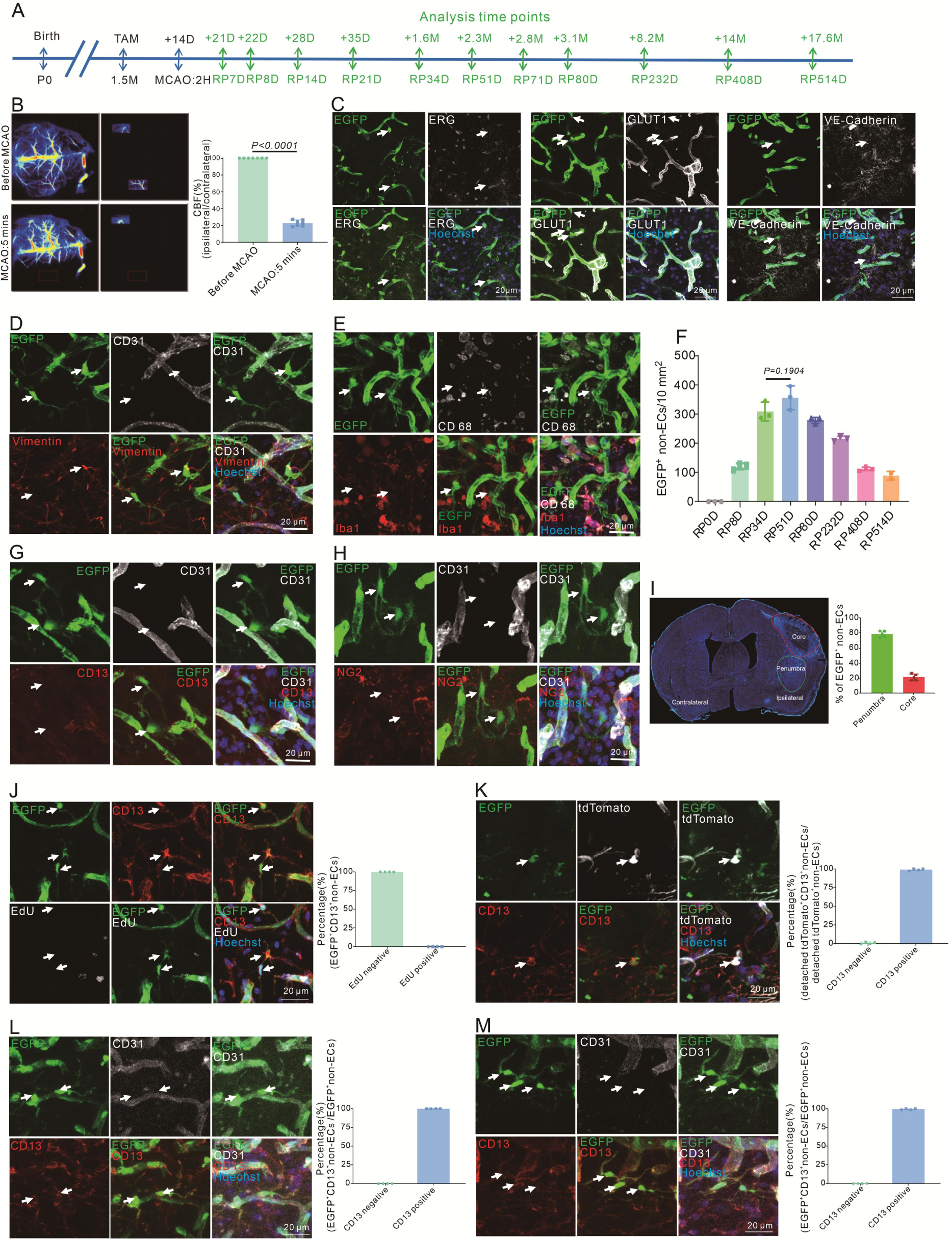
Different means prove that ECs can give rise to pericyte-like cells after stroke, related to. Figure 3. A.Schematic diagram displaying the time course for MCAO, tamoxifen and analysis time points. B.Laser speckle displaying CBF change after stroke(ipsilateral/contralateral) and quantitative the proportion (n=7). C.Immunoflurescence staining of ECs markers (ERG, GLUT1 and VE-Cadherin) expression in Cdh5CreERT2; Ai47 mice with MCAO after RP34D(n=3,20 slices/mouse). D.Immunoflurescence staining of CD31 and vimentin expression in Cdh5CreERT2;Ai47 mice with MCAO at RP34D. E.Immunoflurescence staining of CD68 and Iba1 expression in Cdh5CreERT2;Ai47 mice with MCAO at RP34D. F.Quantitative the number of EGFP^+^&CD31^-^cells in unit area at different reperfusion times after stroke(n=2-6). G.Immunoflurescence staining of CD31 and CD13 expression in Cdh5CreERT2;Ai47 mice with MCAO at RP8D. H.Immunoflurescence staining of CD31 and NG2 expression in Cdh5CreERT2;Ai47 mice with MCAO at RP8D. I.Quantitative the percentage of EGFP^+^&CD31^-^cells in core and penumbra after stroke at RP34D (n=4,20 slices/mouse). J.Immunoflurescence staining of EdU expression in EGFP^+^ pericytes after MCAO at RP34D and quantitative the proportion of EdU^+^ cells(n=4,20 slices/mouse). K.Immunoflurescence staining of CD31 and CD13 expression in Cdh5CreERT2 mice injection with AAV2/9-CAG-DIO-EGFP virus after MCAO at RP34D and quantitative the proportion of CD13^+^ cells(n=4,20 slices/mouse). L.Immunoflurescence staining of CD31 and CD13 expression in Tie2Dre;Mfsd2aXER;Ai47 mice with MCAO at RP34D and quantitative the proportion of CD13^+^ cells(n=4,20 slices/mouse). M.Immunoflurescence staining of CD31 and CD13 expression in Ai47 mice infected with AAV-BI30-Cre virus after MCAO at RP34D and quantitative the proportion of CD13^+^ cells (n=3,20 slices/ mouse).

**Figure S4.**
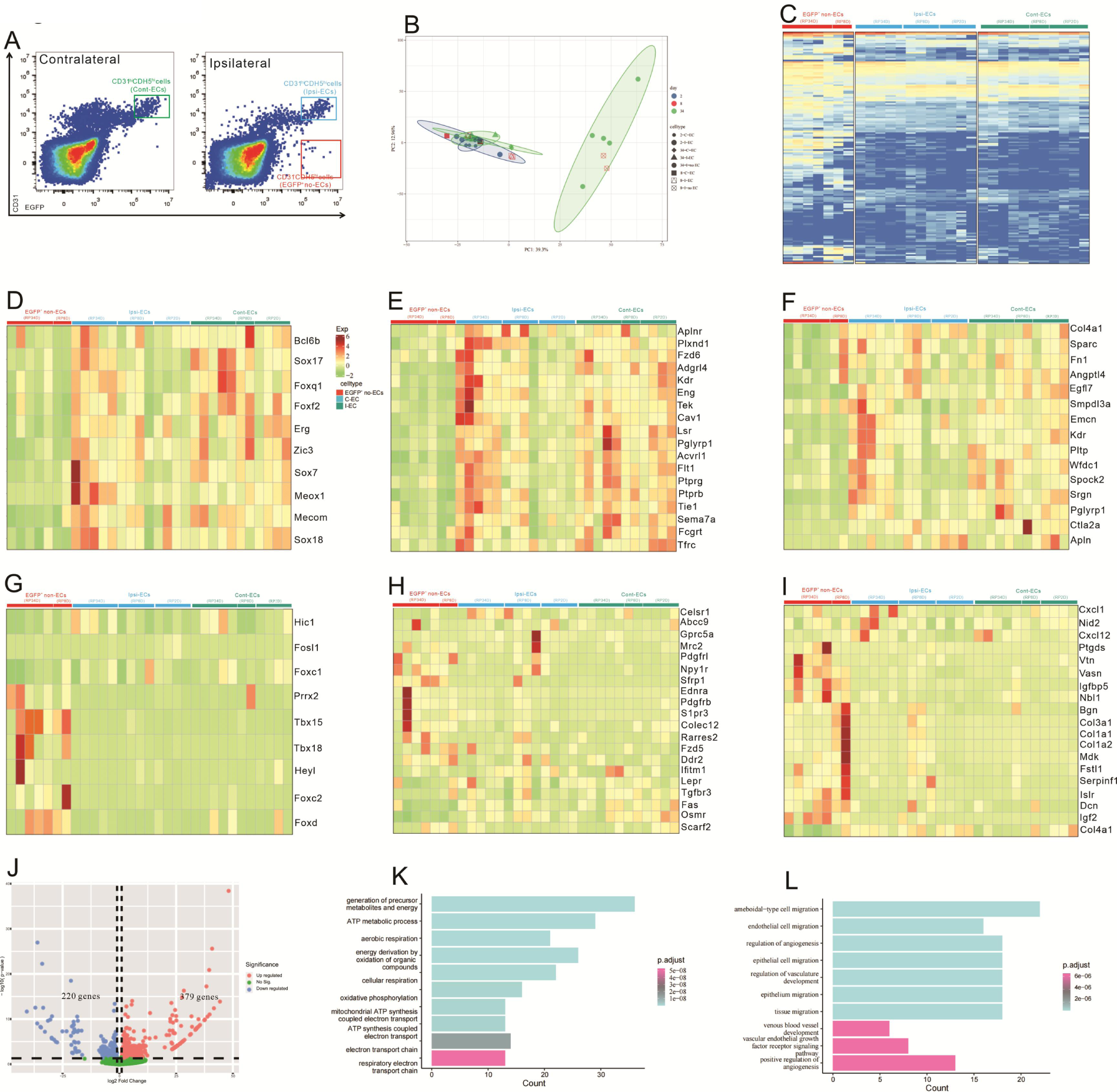
ECs give rise to cells with a similar pericyte transcriptome profile after stroke, related to. Figure 3. A.The gate of sorting ECs in contralateral(C-ECs), ECs in ipsilateral(I-ECs) and EGFP^+^&CD31^-^ cells in ipsilateral(EGFP^+^ non-ECs). B. Principal component analysis (PCA) of the variance-stabilized estimated raw counts of differentially expressed genes. C.The heat map shows 100 genes in all groups. D.The heat map showing endothelial cell transcription factors genes expression in all groups(n=2-5). E.The heat map showing endothelial enriched transmembrane receptor genes expression in all groups(n=2-5). F.The heat map showing endothelial enriched ligand genes expression in all groups(n=2-5). G.The heat map showing pericytic transcription factors genes expression in all groups(n=2-5). H.The heat map showing pericytic-enriched transmembrane receptor gene expression in all groups(n=2-5). I.The heat map showing pericytic enriched ligand genes expression in all groups(n=2-5). J.Volcano plot showing differential expression of genes in EGFP^+^ non-ECs/C-ECs(n=5). K.GO functional enrichment analysis from up-regulation genes in DEG. L.KEGG functional enrichment analysis from up-regulation genes in DEG.

**Figure S5.**
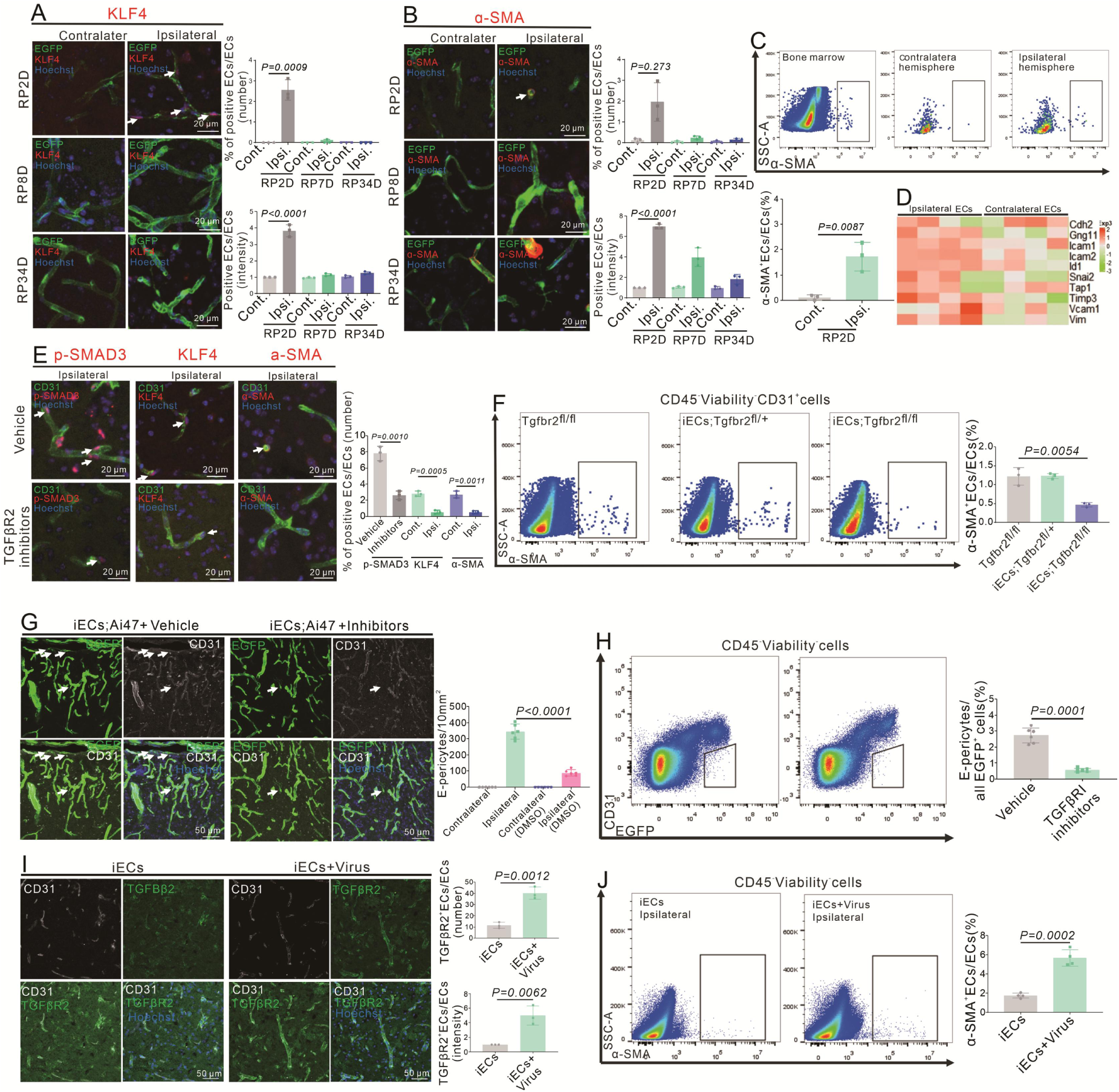
Systemic inhibition of TGFR2 reduces EndoMT and E-Pericytes, related to. Figure 6. A.Immunofluorescence staining of KLF4 expression in Cdh5CreERT2;Ai47 mice at RP2D, RP8D and RP34D and quantitative the proportion and intensity(n=3,20 slices/mouse). B.Immunoflurescence staining of ɑ-SMA expression in Cdh5CreERT2;Ai47 mice and at RP2D, RP8D and RP34D and quantitative the proportion and intensity(n=3,20 slices/mouse). C.Flow cytometry analysis the proportion of ɑ-SMA^+^&CD31^+^ ECs in Cdh5CreERT2; Ai47 mice at RP2D and quantitative the proportion(n=3). D. Heatmap depiction of different EndoMT marker genes in ECs at RP2D(n=4). E.Immunoflurescfence staining of EndoMT markers (p-SMAD3, KLF4 and α-SMA) expression in Cdh5CreERT2; Ai47 mice injection with TGFβR2 inhibitors at RP2D and quantitative the proportion(n=3,20 slices/mouse). F.Flow cytometry analysis the proportion of ɑ-SMA^+^&CD31^+^ ECs in Cdh5CreERT2; Ai47;Tgfbr2^fl/fl^ mice at RP2D and quantitative the proportion (n=3). G.Immunoflurescence staining of CD31 expression in Cdh5CreERT2; Ai47 mice injection with TGFβR2 inhibitors at RP34D and quantitative the number in the unit area (n=3,20 slices/ mouse). H.Flow cytometry analysis of the proportion of E-Pericytes in Cdh5CreERT2; Ai47 mice injection with TGFβR2 inhibitors at RP34D and quantitative the proportion(n=3). I.Immunoflurescence staining of CD31 and TGFBR2 expression in Cdh5CreERT2; Ai47 mice injection with AAV2/9-BI30-EF1α-DIO-Tgfbr2-3XFLAG-P2A-DsRed-WPREs and quantitative the proportion and intensity(n=3,20 slices/mouse). J.Flow cytometry analysis of the proportion of ɑ-SMA+&CD31^+^ ECs in Cdh5CreERT2; Ai47 mice with AAV2/9-BI30-EF1α -DIO-Tgfbr2-3XFLAG-P2A-DsRed-WPREs at RP2D and quantitative the proportion.

**Figure S6.**
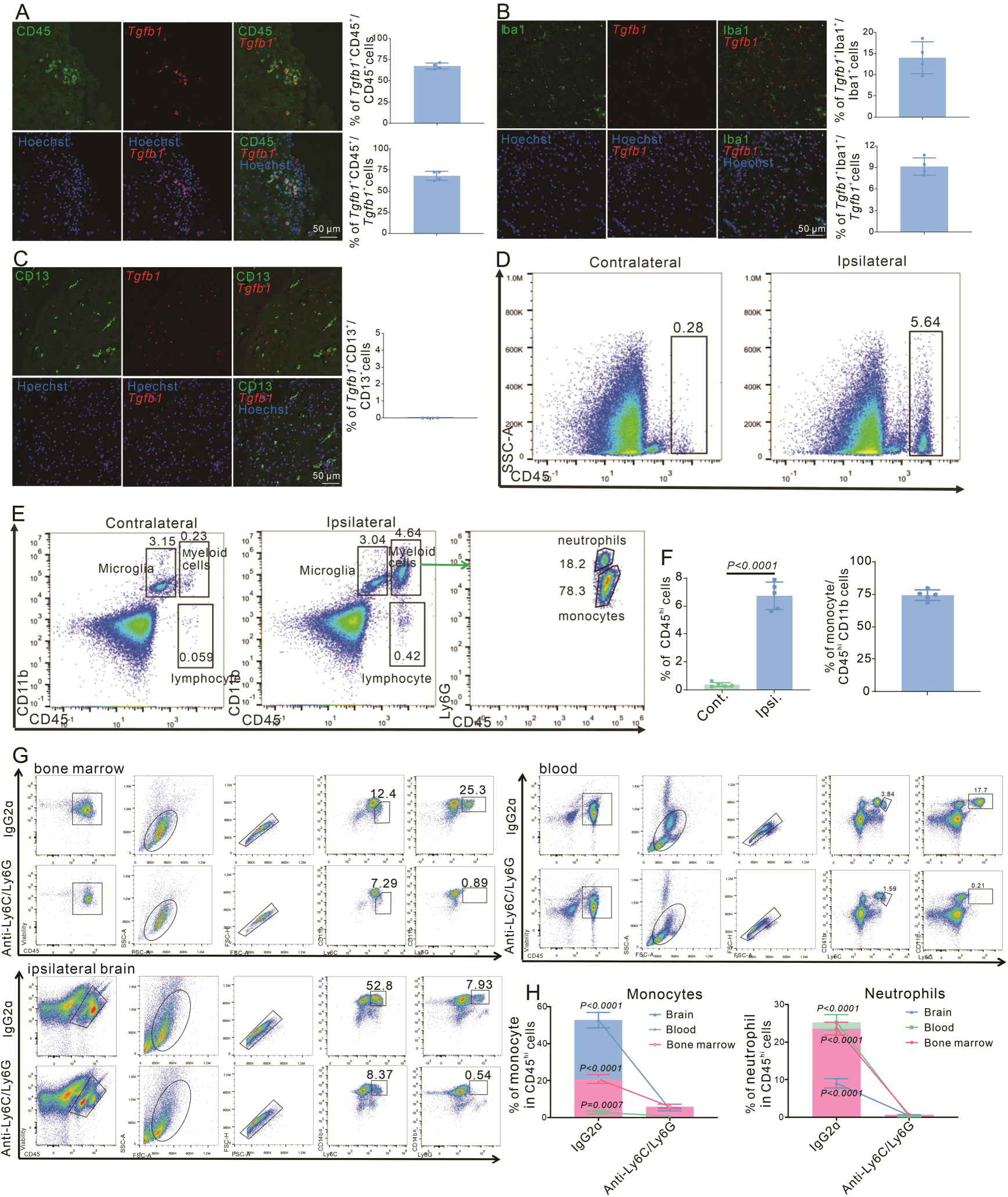
Myeloid cells are leading immune cells in the mouse brain after stroke at RP2D, related to. Figure 7. A.Immunoflurescence staining of CD45 and *Tgfb1* gene expression in WT mice with MCAO at RP1D and quantitative the proportion of *Tgfb1*^+^ cells(n=4,10 slices/mouse). B.Immunoflurescence staining of Iba1 and *Tgfb1* gene expression in WT mice with MCAO at RP2D and quantitative the proportion of *Tgfb1*+ cells(n=4,10 slices/mouse). C.Immunoflurescence staining of CD13 and *Tgfb1* gene expression in WT mice with MCAO at RP2D and quantitative the proportion of *Tgfb1*+ cells(n=4,10 slices/mouse). D.Flow cytometry analysis the proportion of CD45^hi^ cells from ipsilateral brain with MCAO:2H and RP1.5D. E.Flow cytometry analysis the proportion of myeloid cells from the ipsilateral brain with MCAO:2H and RP1.5D. F.Quantitative the proportion of CD45^hi^ cells (D) and monocyte(E)(n=5). G.Flow cytometry analysis the proportion of myeloid cells from bone marrow, blood and ipsilateral brain after mice injection with anti-Ly6C/Ly6G at RP2D(n=4). H. Quantitative the proportion of monocyte and neutrophil in(J)(n=4).

**Figure S7.**
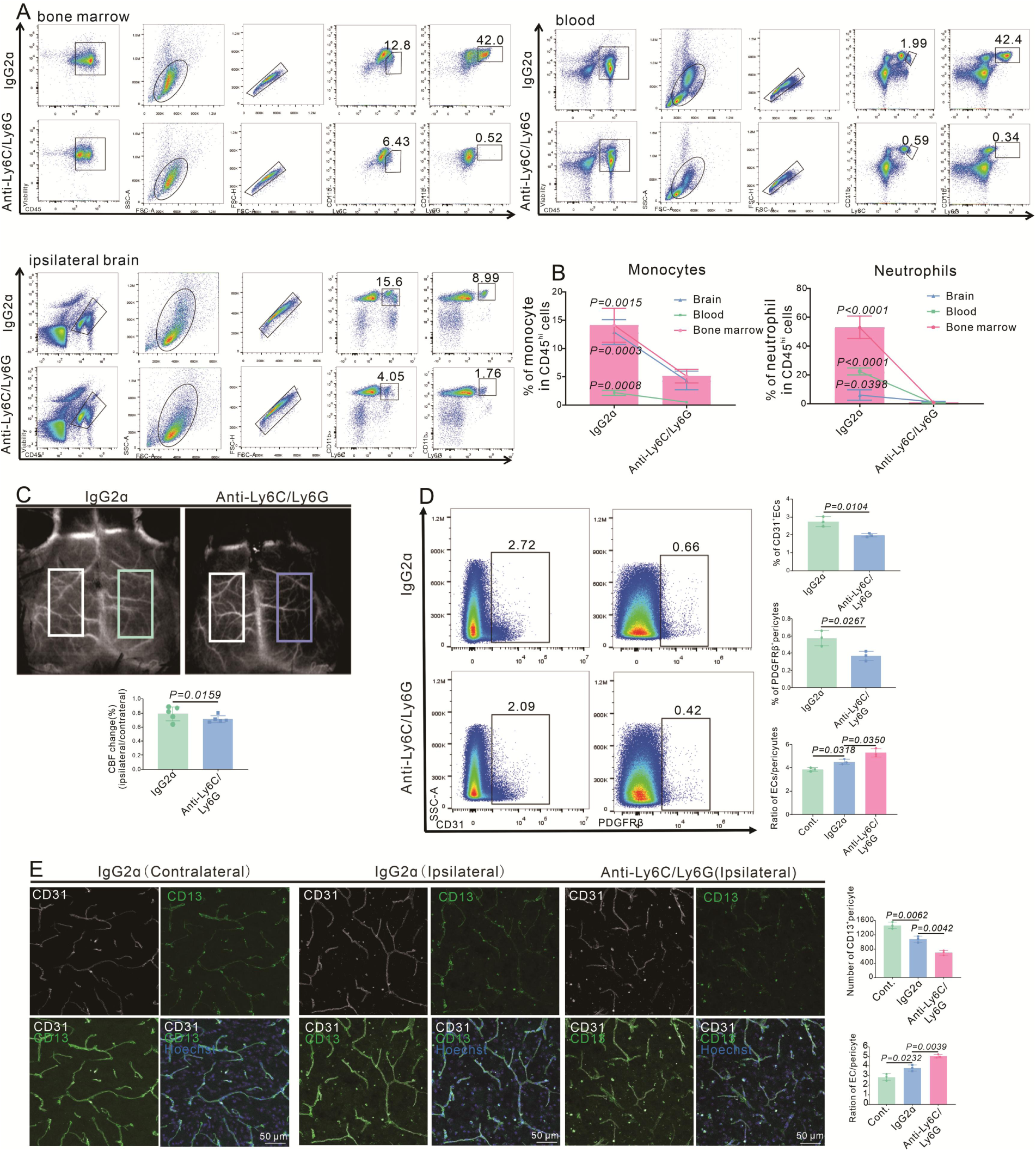
Infiltrating myeloid cells promote vascular function recovery after stroke, related to. Figure 8. A.Flow cytometry analysis the proportion of myeloid cells from bone marrow, blood and ipsilateral brain after mice injection with anti-Ly6C/Ly6G at RP34D(n=4). B.Quantitative the proportion of monocyte and neutrophil in(A)(n=4). C.Image showing the change of CBF in WT mice injection with anti-Ly6C/Ly6G at RP34D after MCAO and quantitative the change of CBF(n=5). D.Flow cytometry analysis the proportion of CD13^+^ cells and CD31^+^ cells in WT mice injection with anti-Ly6C/Ly6G at RP34D after MCAO and quantitative the percentage(n=3). E.Immunofluorescence analysis the proportion of CD13^+^ cells and CD31^+^ cells in WT mice injection with anti-Ly6C/Ly6G at RP34D after MCAO and quantitative the percentage(n=3).

**Figure S8.**
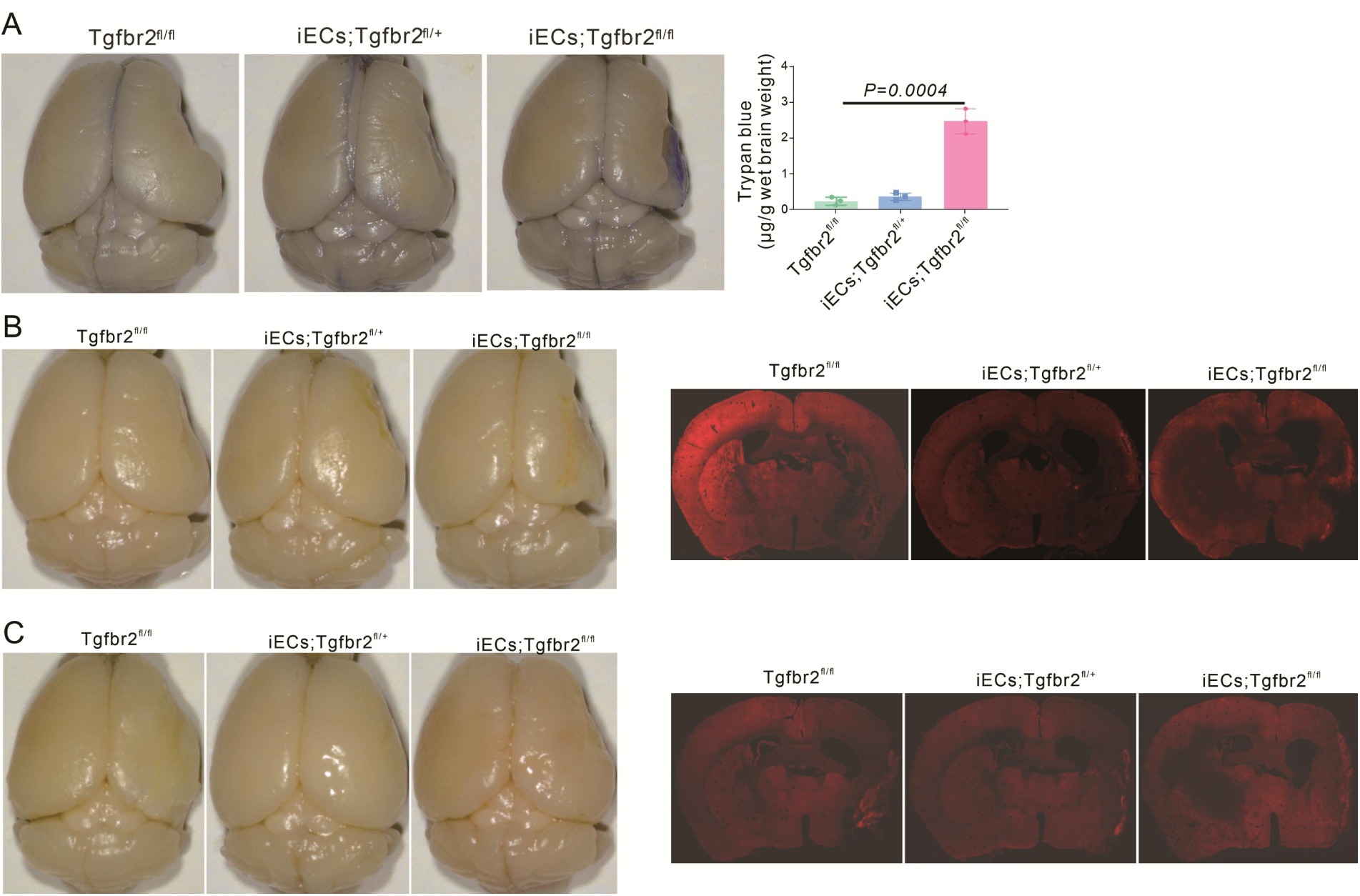
E-Pericytes deletion by specific endothelial cells knockout the *Tgfbr2* gene aggravates BBB leakage small molecule, related to Figure 9. A.Image showing the leakage of Trypan blue in iECs;Tgfbr2^fl/fl^ mice at RP34D after MCAO and quantitative the leakage of Trypan blue(n=3). B.Image showing the leakage of dextran-rhodamine B in iECs;Tgfbr2^fl/fl^ mice at RP34D after MCAO(n=3,10 slices/mouse). C.Image showing the leakage of Texas-Ted in iECs;Tgfbr2^fl/fl^ mice at RP34D after MCAO (n=3,20 slices/mouse).

**Figure S9.**
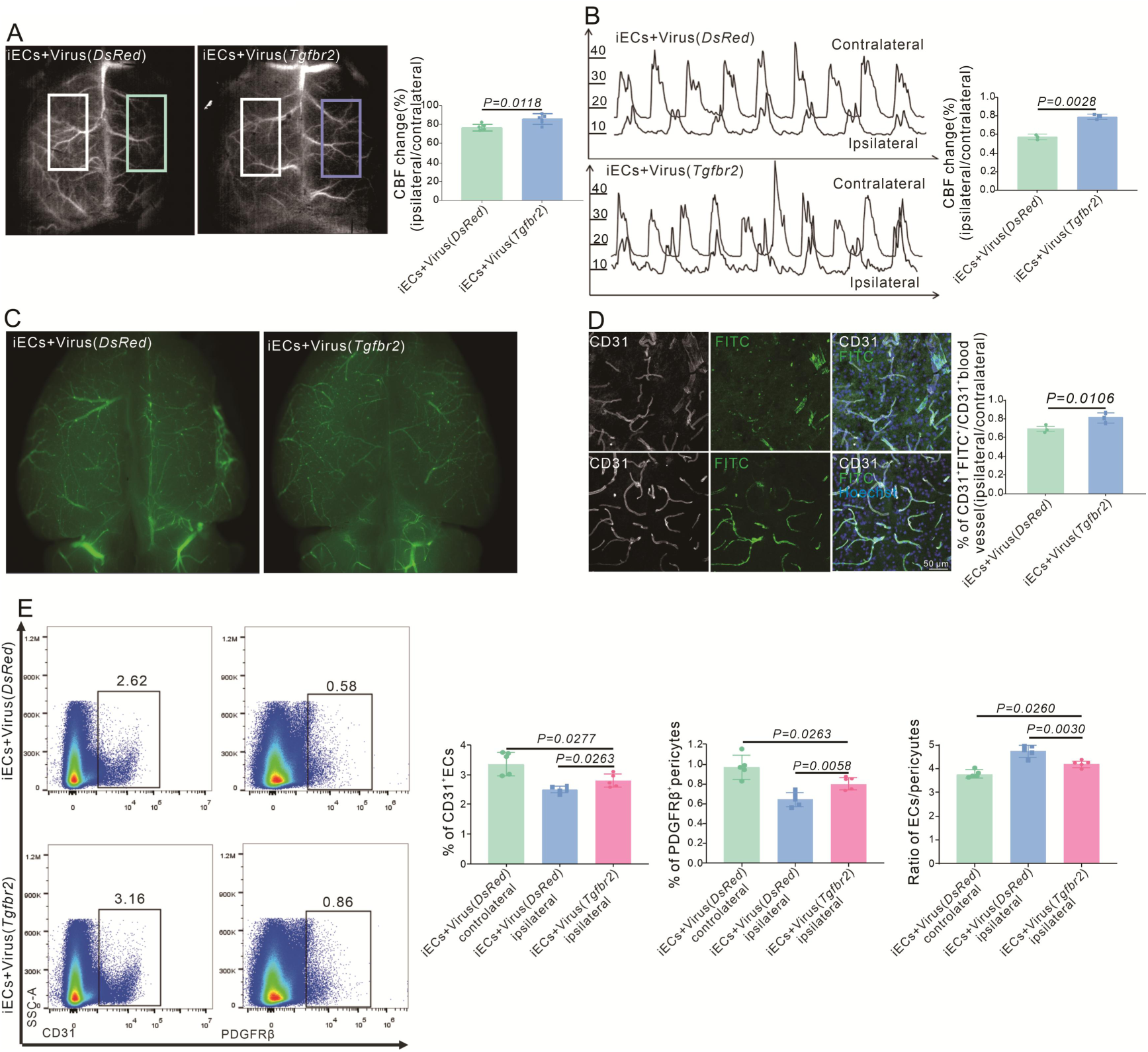
ECs-specific overexpression of the *Tgfbr2* gene reinforces CBF and vessel length, related to. Figure 10. A.Laser speckle image showing the change of CBF in Cdh5CreERT2 mice injection with AAV2/9-BI30-EF1α-DIO-Tgfbr2-3XFLAG-P2A-DsRed-WPREs virus after MCAO at RP34D and quantitative the change of CBF(n=5). B.Graph showing the change of CBF using Doppler test in Cdh5CreERT2 mice injection with AAV2/9-BI30-EF1α-DIO-Tgfbr2-3XFLAG-P2A-DsRed-WPREs virus after MCAO at RP34D and quantitative the change percentage (n=5). C.Image showing FITC-Dextran in gelatin from Cdh5CreERT2 mice injection with AAV2/9-BI30-EF1α-DIO-Tgfbr2-3XFLAG-P2A-DsRed-WPREs virus after MCAO at RP34D and quantitative the percentage(n=3). D.Immunoflurescence staining of CD31 expression in Cdh5CreERT2 mice injection with AAV2/9-BI30-EF1α-DIO-Tgfbr2-3XFLAG-P2A-DsRed-WPREs virus after MCAO at RP34D and quantitative the percentage, FITC-Dextran in gelatin injection by cardiac perfusion (n=3,20 slices/mouse). E.Flow cytometry analysis of the proportion of PDGFR^+^ cells and CD31^+^ cells in Cdh5CreERT2 mice injection with AAV2/9-BI30-EF1α-DIO-Tgfbr2-3XFLAG-P2A-DsRed-WPREs virus after MCAO at RP34D and quantitative the percentage(n=5).

## METHODS

### Experimental design

Transgenic Cdh5CreERT2 transgenic mice cross with reporter mice *Ai47 (EGFP ^fl/fl^) reporter* or *Ai14 (tdTomato^fl/fl^) reporter mice*. Thus, ECs are labeled in green or red initially. TAM was delivered when mice were 1.5 months old (6 weeks), i. e. genetic tracing started. After waiting for 2 weeks, MCAO was performed. After a 2-hour occlusion, reperfusion began. The contralateral sides of the ischemic brain were used as objects for homeostasis conditions. Thus, it was traced for 22 days in the PR8D mice (N = 3 mice), 1.6 months in RP34D (N = 27 mice), 2.8 months in RP71D (N = 3 mice), 8.2 months in RP232D (N = 3 mice), 14 months in RP408D (N = 2 mice),17.6 months in RP51D mice (N = 2 mice).

### Animal care and tissue dissection

All animal experiments were carried out following protocols approved by the Institutional Animal Care and Use Committee (IACUC) at the School of Life Sciences, Westlake University. Wild-type C57BL6/J mice were purchased from the Laboratory Animal Resources Center of Westlake University. Cdh5CreERT2 mice were a gift from Le-Ming Zheng (Peking University). Gt (ROSA)26Sor^tm^^47(CAG-EGFP*)Hze^ (Ai47) was shared by Zilong Qiu (ION). We crossed Cdh5Cre-ERT2 mice to Ai47 mice to generate Cdh5CreERT2:Ai47 mice. Standard chow and water were provided to mice ad libitum. Four mice were housed in each cage. All animals were housed in a standard animal room with a 12/12-hour light/dark cycle at 25 °C. Both male and female mice were used in this study. Mice were anesthetized by intraperitoneal injection of pentobarbital sodium (100mg/kg), and brain dissection was performed immediately for FACS experiments or cardiac PFA perfusion was performed for the immunohistochemistry experiment. After fixation, brains were dissected and kept in 4% PFA for an additional 4 hours and then transferred to PBS until further processing.

### MCAO

Adult mice were anesthetized with pentobarbital sodium (100 mg/kg) and body temperature was maintained during surgery with a heating pad. A midline neck incision was made, the right common carotid artery was carefully separated from the vagus nerve, and the artery was ligated using a 5.0-string. A second knot was made on the left external carotid artery. The right internal carotid artery (ICA) was isolated, and a knot was left loose with a 5.0-string. This knot was not tightened until the intraluminal insertion was done. A small hole was cut in the common carotid artery before it was bifurcated to the external carotid artery and ICA. A silicon-coated monofilament (tip diameter = 230 μm, Doccol Corporation, 602256PK5Re) was gently advanced into the ICA until it stopped at the origin of the MCA in the circle of Willis. The third knot on the ICA was closed to fix the filament in position. During MCA occlusion (2 h), mice were kept in a warm cage at 25 °C.

### Laser speckle contrast imaging (LSCI)

LSCI provides a measure of CBF by quantifying the extent of blurring of dynamic speckles caused by the motion of red blood cells through the vessels. Briefly, mice were placed under an RFLSI III device (RWD Life Sciences) before and after the suture was successfully inserted through the CCA. The skull over both hemispheres was exposed by making an incision along the midline of the scalp. When a 785 nm laser is used to illuminate the brain, it produces a random interference effect that represents blood flow in the form of a speckle pattern. Scattering light was detected by a charge-coupled device (CCD) camera, and the images were acquired by custom software from RWD Life Sciences Company. For each acquisition, a total of 160 images, each of which measured 2048 ×2048 pixels, were collected at 16 Hz.

### Laser Doppler flowmetry (LDF)

After the laser emitted by the LDF device is irradiated onto the tissue, the laser is scattered by the moving red blood cells in the blood, causing a change in the frequency of the scattered light. This frequency change is related to the speed of the red blood cells, so the blood flow velocity can be inferred by measuring the Doppler shift. Briefly, mice were placed under an LDF device (moorVMS-LDF1) before and after the suture was successfully inserted through CCA at RP34D. Connect the probe to the Laser Doppler Flowmeter, turn on the device to start measuring, determine the brain area you wish to measure, position the Laser Doppler probe on the desired brain area, record the data, export the raw data and analysis.

### Tamoxifen

Tamoxifen, 10 mg/ml dissolved in corn oil, was administrated intragastrically with 200ul for 4 consecutive days. Before further experiments, fluorescent protein expressions in mouse ear vasculatures were first examined. Tamoxifen was administrated at least 2 weeks before MCAO surgery.

### Immunohistochemistry

Cryosections of fixed mouse brains (50 μm) were handled free-floating and washed for 5 minutes in PBS before further procedures. Tissue sections were permeablized and blocked in the PBS containing 0.5% Triton and 5% BSA at room temperature for 1 hour. Then, brain sections were incubated with a different primary antibodies, including CD31(1:400, BD, Cat# 557355), VE-Cadherin(1:200, BiCellScientific, Cat# 00105), ERG(1:300, Sigma, Cat# ab110639), GLUT1 (1:300, Sigma, Cat# 07-1401), NeuN (1:300, Merck, Cat# ABN90P), Doublecortin (1:300, Santa clus, Cat# sc-271390), Iba1 (1:300, HUABIO, Cat# ET1705-78), GFAP (1:300, Proteintech, Cat# CL488-60190), PDGFRβ (1:300, ThermoFisher, Cat# 14-1402-82), NG2 (1:300, Merck, Cat# AB5320), CD68 (1:300, BIO-RAD, Cat# MCA1957T), CD45-APC (1:200, Biogend, Cat# 103112), Ki67 (1:300, thermoscientific, Cat# RM-9106-S0), F4/80 (1:300, Cell Signaling, Cat# 30325), CD13 (1:300, R&D system, Cat# AF2335), α-SMA-CY3 (1:300, Sigma, Cat# C6198), which were all diluted in the blocking solution (1% percelin, 5% BSA, and 0.1% Triton in PBS) at 4 °C overnight. After rinsing in PBS 3 times for 5 minutes each, brain slices were incubated with secondary antibodies accordingly, including donkey anti-mouse IgG (A21202, Thermo Fisher Scientific, 1:1,000) and donkey anti-rabbit IgG (A10040, Thermo Fisher Scientific, 1:1,000). They were diluted in the blocking solution. After a 2-hour incubation, brain slices were washed with PBS three times for 15 minutes each. Finally, Hoechst counterstaining was performed for each specimen. After mounting in an anti-fade mounting medium, images were acquired by using a Zeiss LSM800 laser-scanning confocal microscope with a 10X, 20X or 63X objective, with or without Airyscan mode.

### Flow cytometry

Primary cell preparation was conducted as described below. Mouse brains were collected into cold HBSS with glucose and BSA (HBSS, Vendor info) and then were cut into small pieces by sterile scissors, followed by a centrifuge at 1600 rpm/min for 5 mins to get rid of supernatant. The minced tissues after the centrifuge were incubated with collagenase IV (2.5 mg/ml, ROCHE, Cat# 11088858001) at 37 °C with gentle rotation for 30 min in 2 ml of HBSS. Next, samples were passed through a 70-μm filter and washed with HBSS. To obtain higher cell yield, grind the tissue on the 70-μm filter using a syringe piston and guarantee all tissue passes the filter. The following antibodies in 1:200 dilutions were used: CD31-PE-594 (1:200, Biolegend, Cat# 102520), CD45-APC (1:200, Biogend, Cat# 103112), Ly6C-APC (1:200, Biolegend, Cat# 128016), CD11b-PE (1:200, Biolegend, Cat#101212), Viability dye (1:1000, ThermoFisher, Cat#65-0863-14). Antibody incubation was performed on ice for 30 minutes in HBSS with glucose and BSA. All analyses were performed with CytoFLEX LX-5L1(Beckman). FlowJo V10 software was used for data analyses.

### EdU injection

EdU (200 mg/kg, Alfa Aesar Chemical) was injected intraperitoneally for 34 days since MCAO was performed. EdU was initially dissolved in DMSO at the stock concentration of 500mg/ml and was diluted with saline before injection.

### Calculation of brain atrophy volume

Fixed mouse brains were cut with 100 μm and photographed using Axio Zoom.V16. Using ImageJ software, the image is converted to a grayscale image. Click Analyze, then Set Measurements, select Area and select Show Brain Slice Outline. Calculate the area of each brain slice. Cerebral atrophy volume ratio = (sum of white ischemic area of each section) / (sum of brain area of each section) × 100%.

### Calculation of signal length and intensity in immunofluorescence

Fixed mouse brains were cut with 50 μm, stained relevant antibodies by immunofluorescence and photographed using Zeiss LSM 800 Confocal Laser Scanning Microscopy (20X). Using ImageJ software, the single signal in the image is converted to a grayscale image and calculated in length and intensity. The percentage of targeted signal length or intensity/CD31^+^ blood vessel length shows the targeted signal expression in blood vessels.

### Calculation of the number of cells in immunofluorescence

Fixed mouse brains were cut with 50 μm, stained relevant antibodies by immunofluorescence and photographed using Zeiss LSM 800 Confocal Laser Scanning Microscopy (20X, 0.1mm^2^). After maximum intensity projection (MIP), the number of cells was calculated at 0.1mm^2^.

### TGFβR2 inhibitors injection

LY-364947(Sigma, L6293) and SB-431542(SelleckBio, S1067) were dissolved in dimethylsulphoxide (DMSO). For the in vivo inhibition of the TGFβ signaling, LY-364947(10mg/kg) and SB-431542 (10mg/kg) were intraperitoneally injected daily starting MCAO. The control mice were treated in parallel with the vehicle only. The animals were killed at RP34D.

### Single-cell workflow

scRNA-seq was performed using a 10x Next GEM Single-Cell 3’GEM kit v3.1 (10x Genomics) according to the manufacturer’s protocol. Briefly, an equal number of FACS-isolated CD45^-^&Viability^-^&EGFP^+^cells(sorting for ECs) and CD45^+^& Viability^-^ cells(sorting for immune cells) from the brain. ∼1,000 cells per µl and immediately loaded into the 10x Chromium controller. Separate 10x Genomics reactions were used for each group. Generated libraries were sequenced on an Illumina NovaSeq with >27.5 ×10^3^ reads per cell followed by demultiplexing and mapping to the mouse genome (build mm^10^) using CellRanger v5.0 (10x Genomics). Gene expression matrices were generated using the CellRanger software v5.0.1 (10x Genomics) with standard settings and mapping to the mm10 reference mouse genome. The following data analysis was performed using Seurat package v3.0.

### Bulk RNA-seq analysis

Messenger RNAs used for bulk RNA-seq were harvested from the sorted parenchymal EGFP^+^&CD31^+^ cells (contralateral and ipsilateral) and EGFP^+^& CD31^-^ cells(ipsilateral) from adult Cdh5CreERT2:Ai47 mice, which received tamoxifen as adults. RNA was isolated using the RNeasy Plus Mini Kit (Qiagen). RNA-seq libraries were prepared using a TruSeq RNA Library Prep kit v2 (Illumina). The libraries were sequenced on a HiSeq 2500 instrument with single-end 300–400-bp reads (single indexing reads). The normalized gene count matrix for all genes was used as input through the htseq-count script. A heat map was generated using R code to visualize the gene expression levels in different groups.The genes of interest were clustered as neuromediator receptors and ranked according to expression level from high to low. The DESeq2 was used to compare differences in gene expression between different groups.

### *Tgfb1* gene RNAscope

To show which cells synthesize the secreted protein TGFβ1, we used RNAscope to test. Briefly, the brains were fixed in chilled 4% paraformaldehyde (PFA) for 8H at 4 °C, washed twice in 1X PBS, and dehydrated with 15% sucrose and 30% sucrose overnight. Mice’s brains were cut with 14um. The following day, sections were washed twice in 50 μL of 2X saline sodium citrate buffer (SSC) for 10 min. smFISH Probes (Advanced Cell Diagnostics) were pre-heated for 10 minutes at 40 °C, cooled to room temperature (RT) and added to sections to incubate (always in a humidified chamber) for 2 h at 40 °C. Probes were used to target *Tgfb1* mRNA(ACD Cat No. 406201, NM_009370.2). Following probe incubation, sections were washed (always with RNAScope wash buffer for 1 min) four times, incubated in RNAscope AMP-1 solution for 30 minutes at 40 °C, washed four times, incubated in RNAscope AMP-2 solution for 15 min at 40 °C, washed four times, incubated in RNAscope AMP-3 solution for 30 min at 40 °C, washed four times, incubated in RNAscope AMP-4 solution for 15 min at 40 °C and washed four times. For concomitant immunostaining, sections were incubated in 5% BSA blocking buffer for 1 hr at RT, washed once with 1X PBS, and incubated in primary antibody solution [CD31(1:400, BD, Cat# 557355), Iba1 (1:300, HUABIO, Cat# ET1705-78), CD13 (1:300, R&D system, Cat# AF2335),] overnight at 4 °C. The following day, sections were washed three times with 1X PBS for 5 min each and incubated in fluorescently labeled secondary antibody solution diluted 1:1000 in 1X PBS in the dark for 2 h at RT. Sections were washed three times in 1X PBS for 5 min, incubated in 0.5 mg/mL Hoechst 33258 (Sigma-Aldrich) for 2 min at RT, washed once with 1X PBS for 5 min, and mounted on glass slides using PermaFluor (Thermo Fisher). Sections were imaged on a confocal (Zeiss LSM 800).

### Deletion of myeloid cells

For myeloid cells depletion, mice received intraperitoneal injections of 400 µg of anti-Ly6C (InVivoMAb Antibodies, BE0203) and anti-Ly6G (InVivoMAb Antibodies, BP0075) antibodies per mouse starting two days ahead before MCAO, then received intraperitoneal injections of 100 µg at RP0D, RP2D, RP4D, RP6D and RP8D. Myeloid cells were monitored by FACS at RP2D and RP8D to evaluate the functions of anti-Ly6C and anti-Ly6G.

### Measurement of BBB integrity

To evaluate the leakage of Evans blue and Trypan blue, the mice were injected with 100 µL of 2.5% Evans blue (Sigma-Aldrich, St. Louis, MO) and 0.25% Trypan blue (Sigma-Aldrich, 93595) intravenously through the tail vein 1H before fixing using 4% PFA. Then, the brains were removed and imaged under microscopy. The brain was then separated into ipsilateral and contralateral hemispheres. Next, each hemisphere was supplemented with 500 µL of trichloroacetic acid (TCA), transferred to a 55 °C heat block, and incubated for 24 h to extract Evans blue from the tissues. The mixture was centrifuged to pellet any remaining tissue fragments, and absorbance was measured at 610 nm; 500 µL TCAwas used as a blank. The Evans blue extravasated per g tissue was determined. 71KD-Texas Red (10mg/ml, Vector Laboratories, TL-1176) and 70KD-Rhodamine B isothiocyanate-Dextran (10mg/ml, Sigma, R9379) were also injected with 100 µL by tail vein injection, and mice brain fixed with 4% PFA after 1H later.

### Rotarod Test

Rotarod Test Protocol execution by double-blind for Mice to assess their motor coordination and balance. Place the mouse on a rotating rod. The rod gradually accelerates from a low speed to a higher speed (from 4 to 40rpm in 300s). The mouse must maintain balance and motor coordination to stay on the rod. Record the time (latency) it takes for the mouse to fall off the rod. Repeat the test 3 times to get consistent results. Before the test, training Mice for three days, the goal is for mice to be able to walk forward on the rotating rod. Rotarod test recorded on post-stroke 1, 3, 7, 14, 21 and 34 days for 3 times every day with one-hour intervals.

### Rotating beam test (RBT)

The RBT execution by double-blind is used to evaluate the motor, balance and sensory functions of animals. The RBT was performed 3D before the induction of stroke and on post-stroke 1, 3, 7, 14, 21 and 34 days for 3 times every day with one-hour intervals. Briefly, before testing, the mice were subjected to training sessions for three consecutive days. Each training day consisted of three consecutive sessions where the mice traveled across the entire beam. During the test, the mice were placed on a beam rotating at 3 r.p.m., and the fall frequency, average speed and total travel distance were recorded and analyzed.

### Corner test

The corner test execution by double-blind is used to detect unilateral abnormalities of sensory and motor functions in the stroke model. Mouse is placed between two boards each with a dimension of 30×20×1cm^3^. The edges of the two boards are attached at a 30° angle with a small opening along the joint between the two boards to encourage entry into the corner. The mouse is placed between the two angled boards facing the corner and halfway to the corner. The non-ischemic mouse turns either left or right, but the ischemic mouse preferentially turns toward the non-impaired, ipsilateral (right) side. The turns in one versus the other direction are recorded from ten trials for each test. A total of 10 trials are recorded per animal pre-operatively and on indicated days. Corner test recorded on post-stroke 1, 3, 7, 14, 21 and 34 days for 3 times every day at one-hour intervals.

### Adhesive removal test

The adhesive removal test execution by double-blind is widely used in rodents to evaluate sensorimotor dysfunction and motor asymmetry. Briefly, animals are placed into a 15 cm × 25 cm transparent box and two similar adhesive tapes are attached to the hairless part of each forepaw with the same pressure.

The time it takes to contact and remove the stimuli is recorded. In general, animals spend more time contacting and removing the adhesive tape from the contralateral forepaw, while they have no problem contacting and removing the adhesive tape from the ipsilateral forepaw. The adhesive removal test was recorded on post-stroke 1, 3, 7, 14, 21 and 34 days for 3 times every day at one-hour intervals.

### Virus injection

The following adenovirus vector *AAV2/9-CAG-DIO-EGFP* was purchased for ShuMi, Wuhan, China (PT-0168). Half of the one-micron virus was delivered to postnatal day 2 mouse pups by intracerebroventricular injection to both sides. After 1.5 months, tamoxifen was administrated. One month later, these mice were subjected to MCAO to induce ischemic stroke. The following adenovirus vector *AAV2/9-NG2-full length-promoter-DIO-DsRed-WPRES* and *AAV2/9-NG2-full length-promoter-DIO-DTA-WPRES* was purchased for ShuMi, Wuhan, China (PT-9649 and PT-9648). The following adenovirus vector *AAV2/9-BI30-EF1α-DIO-Tgfbr2-3XFLAG-P2A-DsRed-WPREs*{Virus*(Tgfbr2)*} was purchased for ShuMi, Wuhan, China (PT-4190). all virus injection by retro-orbital injection with 1e*10^11^vg.

### Vascular flow function

At RP34D after MCAO, mice were disposed with cardiac perfusion 9 ml/min with 20 ml of warm (34-37°C) PBS with heparin (20 IU/ml), followed by 20 ml of warm (34-37°C) 0.25% (w/v) FITC-Dextran (Sigma-Aldrich, SLCC4853) in 5% (w/v) gelatin from porcine skin (Sigma-Aldrich, G1890) in PBS. After placing the mice heads down into ice for 30 min, the brains were extracted and drop-fixed in 4% PFA overnight for immunofluorescence.

### Mouse cranial imaging by two-photon microscopy

Adult mice were anesthetized with pentobarbital sodium, and analgesia was provided by subcutaneous injection of 0.2% meloxicam. The scalp was removed, and the skull was exposed and cleaned. A dental drill with a 0.6-mm-diameter bit was used to engrave and thin the bone around the circular craniotomy area at a size of 3 mm. The piece of skull was carefully peeled off with fine-pointed forceps, and a cranial window was generated. A coverslip with a 3-mm diameter was placed on the cranial window, and its perimeter was completely sealed with a 1% agarose gel. The metal head plate was glued onto the skull with dental acrylic, through which the mouse brain was fixed on a head plate holder. The headplate holder together with the mouse was placed under a two-photon microscope (FVMPE-RS, Olympus). The cerebral vasculature *Cdh5CreERT2:Ai47* mice were visualized and imaged once a day before and after stroke. All surgeries were performed with sterilized instruments and an environment.

### Statistics

The quantified data in all figures were analyzed with GraphPad Prism 10.0 (La Jolla, CA, USA) and presented as the mean ± SEM with individual data points shown. Unpaired two-tailed Student’s t-test was used to assess the statistical significance between the two groups. Statistical significance was determined by calculation of p-value (*\*p < 0.05, **p < 0.01, ***p < 0.001, and ****p < 0.0001, ns: not significant*). The repetition of our data is independent biological replicates and the number of replicates for each experiment is noted in the corresponding figure legend.

## Acknowledgments

We thank L.He, W.Pei, Z.Gao, H,Shi, Y.Lu, B.Cai and Q.Ma for insightful discussions. We thank Y.Wang for performing part FACS and bulk RNA-seq library preparation together. We thank D.Lu for her responsive and timely purchasing support. We thank Q.Gao for part FACS and mouse genotyping together. We thank Y.Jin for part Stroke Induced with Magnetic Particles (SIMPLE). We thank P.Zhu for a headgear model for two-photon microscope. We thank J.Xie and L.Gao for their Virus(AAV-CAG-DIO-EGFP). We thank the animal facility for its technical assistance with rodent housing, and the Biomedical Research Center platform for technique support. J.-M.J. acknowledges the support from the Key R&D Program of Zhejiang (grant 2024SSYS0031), Zhejiang Province Natural Science Foundation (Project # 2022XHSJJ004), the National Natural Science Foundation of China (Projects # 32170961), HRHI programs 202309002 and 202109013 of Westlake Laboratory of Life Sciences and Biomedicine, Westlake University startup funding, the Westlake Education Foundation. This work is also supported by the National Natural Science Foundation of China (Projects # 82101475) to Z.Z.

## Author contributions

J.-M.J. conceived the research. T.L. and J.-M.J. designed experiments. T.L. performed almost all the experiments and data quantification. L.Y. and N.L. performed endothelial scRNA-seq data analysis. Z.Z. conducted part Bulk RNA-seq analysis and helpful discussions. G.X. performed immune scRNA-seq data analysis. J.T. performed part BBB leakage and behavioristics.

Y.H. performed part behavioristics together. D.Z. and X.L. performed intracerebroventricular injection and retro-orbital injection together with T.L..

L.Z. performed one figure in the result. Y.Z. performed part calculation of blood vessel length. B.Z. performed genotype identification.

## REFERENCES

1. Michalopoulos, G.K., and Bhushan, B. (2021). Liver regeneration: biological and pathological mechanisms and implications. Nat Rev Gastro Hepat 18, 40–55. 10.1038/s41575-020-0342-4.

2. Michalopoulos, G.K., and DeFrances, M.C. (1997). Liver regeneration. Science 276, 60–66. DOI 10.1126/science.276.5309.60.

3. Singhal, M., Liu, X.T., Inverso, D., Jiang, K., Dai, J.N., He, H., Bartels, S., Li, W.P., Pari, A.A.A., Gengenbacher, N., et al. (2018). Endothelial cell fitness dictates the source of regenerating liver vasculature. J Exp Med 215, 2497–2508. 10.1084/jem.20180008.

4. Peña, O.A., and Martin, P. (2024). Cellular and molecular mechanisms of skin wound healing. Nat Rev Mol Cell Bio 25, 599–616. 10.1038/s41580-024-00715-1.

5. Park, S., Gonzalez, D.G., Guirao, B., Boucher, J.D., Cockburn, K., Marsh, E.D., Mesa, K.R., Brown, S., Rompolas, P., Haberman, A.M., et al. (2017). Tissue-scale coordination of cellular behaviour promotes epidermal wound repair in live mice (vol 19, pg 155, 2017). Nat Cell Biol 19, 407–407. DOI 10.1038/ncb3503.

6. Broughton, B.R.S., Reutens, D.C., and Sobey, C.G. (2009). Apoptotic Mechanisms After Cerebral Ischemia. Stroke 40, E331–E339. 10.1161/Strokeaha.108.531632.

7. Gorelick, P.B. (2019). The global burden of stroke: persistent and disabling. Lancet Neurology 18, 417-+. 10.1016/S1474-4422(19)30030-4.

8. Feigin, V.L., Nguyen, G., Cercy, K., Johnson, C.O., Alam, T., Parmar, P.G., Abajobir, A.A., Abate, K.H., Abd-Allah, F., Abejie, A.N., et al. (2018). Global, Regional, and Country-Specific Lifetime Risks of Stroke, 1990 and 2016. New Engl J Med 379, 2429–2437. 10.1056/NEJMoa1804492.

9. Barker, R.A., Gotz, M., and Parmar, M. (2018). New approaches for brain repair-from rescue to reprogramming. Nature 557, 329–334. 10.1038/s41586-018-0087-1.

10. Gill, S.S., Patel, N.K., Hotton, G.R., O’Sullivan, K., McCarter, R., Bunnage, M., Brooks, D.J., Svendsen, C.N., and Heywood, P. (2003). Direct brain infusion of glial cell line-derived neurotrophic factor in Parkinson disease. Nat Med 9, 589–595. 10.1038/nm850.

11. Bartus, R.T., and Johnson, E.M. (2017). Clinical tests of neurotrophic factors for human neurodegenerative diseases, part 1: Where have we been and what have we learned? Neurobiol Dis 97, 156–168. 10.1016/j.nbd.2016.03.027.

12. Bartus, R.T., and Johnson, E.M. (2017). Clinical tests of neurotrophic factors for human neurodegenerative diseases, part 2: Where do we stand and where must we go next? Neurobiol Dis 97, 169–178. 10.1016/j.nbd.2016.03.026.

13. Juttler, E., Kohrmann, M., and Schellinger, P.D. (2006). Therapy for early reperfusion after stroke. Nat Clin Pract Card 3, 656–663. 10.1038/ncpcardio0721.

14. Li, S.Y., Gu, H.Q., Li, H., Wang, X.C., Jin, A.M., Guo, S.M., Lu, G.Z., Che, F.Y., Wang, W.W., Wei, Y., et al. (2024). Reteplase versus Alteplase for Acute Ischemic Stroke. New Engl J Med 390, 2264–2273. 10.1056/NEJMoa2400314.

15. Murrant, C.L., and Fletcher, N.M. (2022). Capillary communication: the role of capillaries in sensing the tissue environment, coordinating the microvascular, and controlling blood flow. Am J Physiol-Heart C 323, H1019–H1036. 10.1152/ajpheart.00088.2022.

16. Iadecola, C., Smith, E.E., Anrather, J., Gu, C.H., Mishra, A., Misra, S., Perez-Pinzon, M.A., Shih, A.Y., Sorond, F.A., van Veluw, S.J., et al. (2023). The Neurovasculome: Key Roles in Brain Health and Cognitive Impairment: A Scientific Statement From the American Heart Association/American Stroke Association. Stroke 54, E251–E271. 10.1161/Str.0000000000000431.

17. Trimm, E., and Red-Horse, K. (2023). Vascular endothelial cell development and diversity. Nat Rev Cardiol 20, 197–210. 10.1038/s41569-022-00770-1.

18. Hall, C.N., Reynell, C., Gesslein, B., Hamilton, N.B., Mishra, A., Sutherland, B.A., O’Farrell, F.M., Buchan, A.M., Lauritzen, M., and Attwell, D. (2014). Capillary pericytes regulate cerebral blood flow in health and disease. Nature 508, 55–60. 10.1038/nature13165.

19. Ding, J., Lee, S.J., Vlahos, L., Yuki, K., Rada, C.C., van Unen, V., Vuppalapaty, M., Chen, H., Sura, A., McCormick, A.K., et al. (2023). Therapeutic blood-brain barrier modulation and stroke treatment by a bioengineered FZD-selective WNT surrogate in mice. Nat Commun 14. ARTN 294710.1038/s41467-023-37689-1.

20. Armulik, A., Genove, G., Mae, M., Nisancioglu, M.H., Wallgard, E., Niaudet, C., He, L., Norlin, J., Lindblom, P., Strittmatter, K., et al. (2010). Pericytes regulate the blood-brain barrier. Nature 468, 557–561. 10.1038/nature09522.

21. Yokomizo, T., and Suda, T. (2024). Development of the hematopoietic system: expanding the concept of hematopoietic stem cell-independent hematopoiesis. Trends Cell Biol 34, 161–172. 10.1016/j.tcb.2023.06.007.

22. Xu, Y., and Kovacic, J.C. (2023). Endothelial to Mesenchymal Transition in Health and Disease. Annu Rev Physiol 85, 245–267. 10.1146/annurev-physiol-032222-080806.

23. Chen, Q., Zhang, H., Liu, Y., Adams, S., Eilken, H., Stehling, M., Corada, M., Dejana, E., Zhou, B., and Adams, R.H. (2016). Endothelial cells are progenitors of cardiac pericytes and vascular smooth muscle cells. Nat Commun 7. ARTN 1242210.1038/ncomms12422.

24. Yu, Q.C., Song, W.Q., Wang, D.S., and Zeng, Y.A. (2016). Identification of blood vascular endothelial stem cells by the expression of protein C receptor. Cell Res 26, 1079–1098. 10.1038/cr.2016.85.

25. Jiang, D.Y., and Mccullough, L. (2024). Mapping brain-immune interactions in ischemic stroke. Nat Immunol 25, 396–398. 10.1038/s41590-024-01747-7.

26. Beuker, C., Schafflick, D., Strecker, J.K., Heming, M., Li, X.L., Wolbert, J., Schmidt-Pogoda, A., Thomas, C., Kuhlmann, T., Aranda-Pardos, I., et al. (2022). Stroke induces disease-specific myeloid cells in the brain parenchyma and pia. Nat Commun 13. ARTN 94510.1038/s41467-022-28593-1.

27. Shichita, T., Ooboshi, H., and Yoshimura, A. (2023). Neuroimmune mechanisms and therapies mediating post-ischaemic brain injury and repair. Nature Reviews Neuroscience 24, 299–312. 10.1038/s41583-023-00690-0.

28. Gliem, M., Mausberg, A.K., Lee, J.I., Simiantonakis, I., van Rooijen, N., Hartung, H.P., and Jander, S. (2012). Macrophages prevent hemorrhagic infarct transformation in murine stroke models. Ann Neurol 71, 743–752. 10.1002/ana.23529.

29. Sas, A.R., Carbajal, K.S., Jerome, A.D., Menon, R., Yoon, C., Kalinski, A.L., Giger, R.J., and Segal, B.M. (2020). A new neutrophil subset promotes CNS neuron survival and axon regeneration. Nat Immunol 21, 1496-+. 10.1038/s41590-020-00813-0.

30. Fang, W.R., Zhai, X., Han, D., Xiong, X.X., Wang, T., Zeng, X., He, S.C., Liu, R., Miyata, M., Xu, B.H., and Zhao, H. (2018). CCR2-dependent monocytes/macrophages exacerbate acute brain injury but promote functional recovery after ischemic stroke in mice. Theranostics 8, 3530–3543. 10.7150/thno.24475.

31. Pedragosa, J., Mir-Mur, F., Otxoa-de-Amezaga, A., Justicia, C., Ruz-Jan, F., Ponsaerts, P., Pasparakis, M., and Planas, A.M. (2020). CCR2 deficiency in monocytes impairs angiogenesis and functional recovery after ischemic stroke in mice. J Cerebr Blood F Met 40, S98–S116. Artn 0271678x2090905510.1177/0271678x20909055.

32. Wattananit, S., Tornero, D., Graubardt, N., Memanishvili, T., Monni, E., Tatarishvili, J., Miskinyte, G., Ge, R.M., Ahlenius, H., Lindvall, O., et al. (2016). Monocyte-Derived Macrophages Contribute to Spontaneous Long-Term Functional Recovery after Stroke in Mice. J Neurosci 36, 4182–4195. 10.1523/Jneurosci.4317-15.2016.

33. Doyle, K.P., Cekanaviciute, E., Mamer, L.E., and Buckwalter, M.S. (2010). TGFβ signaling in the brain increases with aging and signals to astrocytes and innate immune cells in the weeks after stroke. J Neuroinflamm 7. Artn 6210.1186/1742-2094-7-62.

34. Krupinski, J., Kumar, P., Kumar, S., and Kaluza, J. (1996). Increased expression of TGF-beta 1 in brain tissue after ischemic stroke in humans. Stroke 27, 852–857. Doi 10.1161/01.Str.27.5.852.

35. Cooley, B.C., Nevado, J., Mellad, J., Yang, D., St Hilaire, C., Negro, A., Fang, F., Chen, G.B., San, H., Walts, A.D., et al. (2014). TGF-β Signaling Mediates Endothelial-to-Mesenchymal Transition (EndMT) During Vein Graft Remodeling. Sci Transl Med 6. ARTN 227ra3410.1126/scitranslmed.3006927.

36. Payne, S., De Val, S., and Neal, A. (2018). Endothelial-Specific Cre Mouse Models: Is Your Cre CREdibile? Arterioscl Throm Vas 38, 2550–2561. 10.1161/Atvbaha.118.309669.

37. Vanlandewijck, M., He, L.Q., Mäe, M.A., Andrae, J., Ando, K., Del Gaudio, F., Nahar, K., Lebouvier, T., Laviña, B., Gouveia, L., et al. (2018). A molecular atlas of cell types and zonation in the brain vasculature (vol 554, pg 475, 2018). Nature 560. 10.1038/s41586-018-0232-x.

38. Kalucka, J., de Rooij, L.P.M.H., Goveia, J., Rohlenova, K., Dumas, S.J., Meta, E., Conchinha, N.V., Taverna, F., Teuwen, L.A., Veys, K., et al. (2020). Single-Cell Transcriptome Atlas of Murine Endothelial Cells. Cell 180, 764-+. 10.1016/j.cell.2020.01.015.

39. Garcia-Bonilla, L., Shahanoor, Z., Sciortino, R., Nazarzoda, O., Racchumi, G., Iadecola, C., and Anrather, J. (2024). Analysis of brain and blood single-cell transcriptomics in acute and subacute phases after experimental stroke. Nat Immunol 25. 10.1038/s41590-023-01711-x.

40. Zhang, Y., Chen, K.N., Sloan, S.A., Bennett, M.L., Scholze, A.R., O’Keeffe, S., Phatnani, H.P., Guarnieri, P., Caneda, C., Ruderisch, N., et al. (2015). An RNA-Sequencing Transcriptome and Splicing Database of Glia, Neurons, and Vascular Cells of the Cerebral Cortex (vol 35, pg 11929, 2014). J Neurosci 35, 864–866. 10.1523/Jneurosci.4506-14.2015.

41. Teng, Z.Q., Ma, Y.T., and Ma, X. (2024). Cardiac Pericytes Acquire a Fibrogenic Phenotype and Contribute to Vascular Maturation After Myocardial Infarction. Circulation 149, e960–e961. 10.1161/Circulationaha.123.066563.

42. Gentek, R., Ghigo, C., Hoeffel, G., Bulle, M.J., Msallam, R., Gautier, G., Launay, P., Chen, J.M., Ginhoux, F., and Bajénoff, M. (2018). Hemogenic Endothelial Fate Mapping Reveals Dual Developmental Origin of Mast Cells. Immunity 48, 1160-+. 10.1016/j.immuni.2018.04.025.

43. Prazeres, P.H.D.M., Almeida, V.M., Lousado, L., Andreotti, J.P., Paiva, A.E., Santos, G.S.P., Azevedo, P.O., Souto, L., Almeida, G.G., Filev, R., et al. (2018). Macrophages Generate Pericytes in the Developing Brain. Cell Mol Neurobiol 38, 777–782. 10.1007/s10571-017-0549-2.

44. De Bock, K., Georgiadou, M., Schoors, S., Kuchnio, A., Wong, B.W., Cantelmo, A.R., Quaegebeur, A., Ghesquière, B., Cauwenberghs, S., Eelen, G., et al. (2013). Role of PFKFB3-Driven Glycolysis in Vessel Sprouting. Cell 154, 651–663. 10.1016/j.cell.2013.06.037.

45. Falkenberg, K.D., Rohlenova, K., Luo, Y.L., and Carmeliet, P. (2019). The metabolic engine of endothelial cells. Nat Metab 1, 937–946. 10.1038/s42255-019-0117-9.

46. Ayloo, S., Lazo, C.G., Sun, S.H., Zhang, W., Cui, B.X., and Gu, C.H. (2022). Pericyte-to-endothelial cell signaling via vitronectin-integrin regulates blood-CNS barrier. Neuron 110, 1641-+. 10.1016/j.neuron.2022.02.017.

47. Andjelkovic, A.V., Situ, M., Citalan-Madrid, A.F., Stamatovic, S.M., Xiang, J.M., and Keep, R.F. (2023). Blood-Brain Barrier Dysfunction in Normal Aging and Neurodegeneration: Mechanisms, Impact, and Treatments. Stroke 54, 661–672. 10.1161/Strokeaha.122.040578.

48. Zhao, Z., Nelson, A.R., Betsholtz, C., and Zlokovic, B.V. (2015). Establishment and Dysfunction of the Blood-Brain Barrier. Cell 163, 1064–1078. 10.1016/j.cell.2015.10.067.

49. Li, Y., Lui, K.O., and Zhou, B. (2018). Reassessing endothelial-to-mesenchymal transition in cardiovascular diseases. Nat Rev Cardiol 15, 445–456. 10.1038/s41569-018-0023-y.

50. Jenkins, G. (2008). The role of proteases in transforming growth factor-β activation. Int J Biochem Cell B 40, 1068–1078. 10.1016/j.biocel.2007.11.026.

51. Kratofil, R.M., Shim, H.B., Shim, R., Lee, W.Y., Labit, E., Sinha, S., Keenan, C.M., Surewaard, B.G.J., Noh, J.Y., Sun, Y.X., et al. (2022). A monocyte-leptin-angiogenesis pathway critical for repair post-infection. Nature 609, 166-+. 10.1038/s41586-022-05044-x.

52. Chi, Z.X., Chen, S., Yang, D.H., Cui, W.Y., Lu, Y., Wang, Z., Li, M.B., Yu, W.W., Zhang, J., Jiang, Y., et al. (2024). Gasdermin D-mediated metabolic crosstalk promotes tissue repair. Nature 634. 10.1038/s41586-024-08022-7.

53. Hall, C.N., Reynell, C., Gesslein, B., Hamilton, N.B., Mishra, A., Sutherland, B.A., O’Farrell, F.M., Buchan, A.M., Lauritzen, M., and Attwell, D. (2014). Capillary pericytes regulate cerebral blood flow in health and disease. Nature 508, 55-+. 10.1038/nature13165.

54. Svara, F., Förster, D., Kubo, F., Januszewski, M., dal Maschio, M., Schubert, P.J., Kornfeld, J., Wanner, A.A., Laurell, E., Denk, W., and Baier, H. (2022). Automated synapse-level reconstruction of neural circuits in the larval zebrafish brain. Nat Methods 19, 1357-+. 10.1038/s41592-022-01621-0.

55. Sanes, J.R., and Zipursky, S.L. (2020). Synaptic Specificity, Recognition Molecules, and Assembly of Neural Circuits (vol 181, pg 536, 2020). Cell 181, 1434–1435. 10.1016/j.cell.2020.05.046.

56. Goumans, M.J., Liu, Z., and ten Dijke, P. (2009). TGF-β signaling in vascular biology and dysfunction. Cell Res 19, 116–127. 10.1038/cr.2008.326.

57. ten Dijke, P., and Arthur, H.M. (2007). Extracellular control of TGFβ signaling in vascular development and disease. Nat Rev Mol Cell Bio 8, 857–869. 10.1038/nrm2262.

